# A mouse-adapted *Staphylococcus aureus* strain enables lifelong neonatal colonization and elicits a Th17-dominated immune response

**DOI:** 10.64898/2026.08.28.747726

**Authors:** Liliane Maria Fernandes Hartzig, Sean Lando Levin Seegert, Murthy Narayana Darisipudi, Shruthi Peringathara, Frieder Schmiedeke, Daniel M. Mrochen, Stefan Gross, Stefan Weiss, Barbara M. Bröker, Silva Holtfreter

## Abstract

The opportunistic pathogen *Staphylococcus aureus* persistently colonizes the anterior nares of up to 20% of the human population, yet there were no persistent mouse colonization models to study host-pathogen interaction. Using the mouse-adapted *S. aureus* strain JSNZ (CC88-MSSA), we established a neonatal *S. aureus* colonization model in C57BL/6N mice. Natural neonatal colonization was achieved by vertical transmission in a JSNZ-positive breeding colony. Offspring were followed for up to 69 weeks and found persistently colonized in the nose and cecum with high bacterial loads. Adult mice were colonized by intranasal inoculation of JSNZ; controls received PBS. The colonization patterns and the *S. aureus*-specific T cell responses were then monitored over a period of 28 days and compared between age-matched mice colonized as neonates or adults. The neonatal group remained persistently colonized in nose and gut with high bacterial densities. In contrast, mice colonized as adults had lower and declining bacterial loads in the nose. Some eliminated *S. aureus* from the nares, while all remained colonized in the gut. Neonatally colonized mice exhibited reduced nasal chemokine levels, which may have favored the prolonged *S. aureus* persistence. *Ex vivo* re-stimulation of cervical lymph node cells with an *S. aureus* antigen cocktail revealed a Th17-dominated antigen-specific T cell response in both colonized groups. The lymph node cells secreted large amounts of IL-17, but Th1-, Th2-associated and regulatory cytokines were also detected. The cytokine patterns were similar in both colonized groups except for IL-5, which was more abundant upon neonatal colonization. In conclusion, vertical transmission of the mouse-adapted *S. aureus* strain JSNZ reliably establishes persistent high-density neonatal colonization, providing a physiologically relevant model for the study of *S. aureus* host interactions. Route and timing of colonization do not fundamentally affect the T cell response to *S. aureus*.

**Author summary:** Up to 20% of the population carry the bacterium *Staphylococcus aureus* in their nose for months or years, usually without symptoms. When the immune system is weakened, however, these bacteria can cause serious infections. Therefore, it is important to understand how our immune system recognizes *S. aureus* during colonization — and how the bacterium persists despite this.

Research in this area has long been hampered by the lack of animal models that reliably reproduce long-term colonization. We addressed this using a mouse-adapted *S. aureus* strain, JSNZ, which colonized adult mice for several weeks. Colonized parents transmitted the strain to their neonates who then carry the bacteria for their lifetime. They had reduced nasal levels of immune-cell recruiting proteins (chemokines), which may have contributed to prolonged bacterial persistence. Both neonatal and adult colonization triggered a strong, lasting immune response dominated by Th17 cells. This response was comparable regardless of whether colonization occurred in early life or adulthood.

This model offers a physiologically relevant tool to study *S. aureus*-immune system interactions during colonization, advancing our understanding of infection risk and potential preventive strategies.

## Introduction

*Staphylococcus (S.) aureus* is an opportunistic pathogen that persistently colonizes the nose of approximately 20% of the human population, with the remainder being intermittently colonized by different strains [1,2]. Although usually asymptomatic, colonization with *S. aureus* is a risk factor for life-threatening endogenous infections, including bacteremia and endocarditis [3,4]. The treatment of staphylococcal infections is often hampered by multiple antibiotic resistances [5], and so far, all efforts to develop an anti-*S. aureus* vaccine have failed [6]. A major obstacle to successful vaccine development has been the lack of translation from mouse models to humans in clinical trials [6,7]. For instance, persistent nasal *S. aureus* carriage could not be mimicked in animal models, and our understanding of the host pathogen interaction and adaptive immune response to persistent colonization is still incomplete [8].

Beyond the anterior nares, *S. aureus* colonizes the throat, skin, intestine, and other body sites [9–11]. Bacterial acquisition and persistence are shaped by host immunity, microbiome composition, age, and environment [2,10,11]. Notably, neonates are disproportionately susceptible: prevalence of nasal carriage reaches circa 50% in newborns, declining sharply in the first year of life [12,13], and also intestinal colonization is far more common in neonates than adults [14]. Infants born to colonized mothers are at particularly elevated risk of colonization [13,15]. This high susceptibility in neonates likely reflects limited competition by the still diversifying microbiota and the tolerogenic status of the developing neonatal immune system [10,16–20].

T cells are central to restricting nasal *S. aureus* colonization, with Th17 cells playing a critical role. In mice, clearance of colonization depends on T cells and neutrophils [21]. IL-17A/F knockout mice fail to clear nasal *S. aureus*, as Th17-derived IL-17 drives antimicrobial peptide expression and neutrophil recruitment [21,22]. Conversely, IL-10-mediated suppression of local IL-17-and IL-22-secreting T cells promotes persistence [23]. In humans, circulating *S. aureus*-specific T cells are dominated by Th17 and Th1 cells, with a smaller Treg component [24–27]. Studying these responses is complicated by superantigens and pore-forming toxins that interfere with functional assays [28], an obstacle that can be circumvented by using heat-inactivated superantigen-negative bacteria or recombinant proteins as recall antigens [24–27]. The *S. aureus*-specific T cell response to persistent murine colonization, however, is still unknown.

Persistent *S. aureus* carriage can be achieved in mice by using mouse-adapted *S. aureus* strains. Our group has shown that a substantial proportion of SPF laboratory mice are naturally *S. aureus*-colonized [29–31]. Mouse-adapted strains such as JSNZ (CC88-MSSA) have co-evolved with murine hosts, conferring stable long-term colonization without antibiotic pretreatment — unlike human-adapted strains [29,32]. These strains are therefore valuable tools for studying drug and vaccine efficacy [33,34], bacterial pathogenesis [32,35–37], and anti-*S. aureus* immunity [32].

Here, we exploited natural vertical transmission of JSNZ from colonized breeding pairs to establish a persistent neonatal colonization model [31]. We employed this model to compare colonization dynamics and *S. aureus*-specific T cell responses between mice colonized as neonates or adults.

## Results

### Vertical transmission from *S. aureus*-colonized breeders induces natural persistent neonatal colonization

*S. aureus* colonization in human infants is common, with the mother often being the source of the colonizing strain [13]. However, the lack of suitable animal models has hampered research into the causes and consequences of neonatal transmission. To address this, we employed the mouse-adapted strain JSNZ to establish a *S. aureus*-positive C57BL/6N breeding colony. Adult male and female mice were inoculated intranasally with JSNZ and mated 7 days later. All breeding pairs remained persistently colonized in the nose throughout the entire breeding period (18-47 weeks) (Fig 1A). Males exhibited stable fecal bacterial loads over time, while females showed more variable colonization densities.

**Fig 1.**
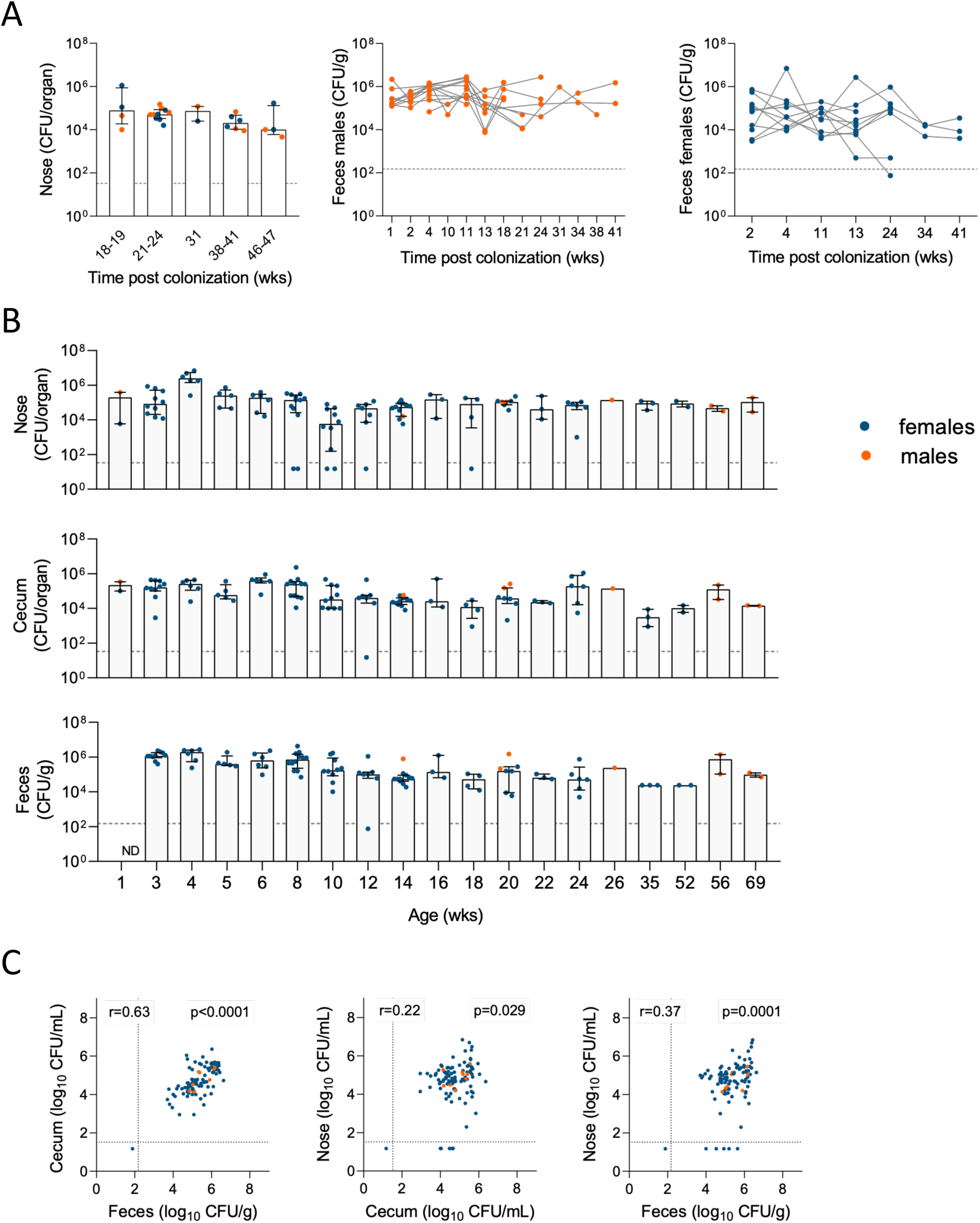
Vertical transmission from *S. aureus*-colonized breeding pairs induces persistent neonatal colonization. Female and male C57BL/6N mice were inoculated intranasally (i.n.) with 1 × 10⁸ CFU *S. aureus* JSNZ and mated 7 days later. (A, B) Bacterial loads in the nose, cecum, and feces of breeding pairs (A) and offspring (B) were determined at the indicated time points post-colonization or post-birth by homogenization and plating on selective agar. (C) Spearman correlation of nasal, cecal and fecal bacterial loads in offspring. Dashed lines indicate the lower limit of detection (LLOD), negative samples are depicted as LLOD/2. Data (A, B) are presented as median with interquartile range (IQR). CFU, colony-forming units; ND, not determined.

Vertical transmission induced persistent nasal and gastrointestinal colonization with JSNZ in the offspring for more than a year (Fig 1B). In only 5 of 102 offspring JSNZ was undetectable in the nose but detectable in the gut at the time of sampling (between 8 - 18 weeks of age). Complete eradication of *S. aureus* was observed in only a single animal. Bacterial loads were consistently high in the nose, cecum, and feces (log₁₀ CFU, mean ± SD: nose 4.67 ± 1.12, cecum 4.75 ± 0.78, feces 4.63 ± 0.77 CFU/organ or ml). As expected, cecal and fecal loads were moderately correlated (r = 0.64) (Fig 1C). Colonization was asymptomatic in females; however, more than 65% of colonized males developed endogenous preputial gland infection (preputial gland adenitis), described in detail in Fernandes Hartzig et al. [38]. Together, these results show that persistent neonatal *S. aureus* colonization can be efficiently established by vertical transmission without further manipulation of the offspring.

### Neonatal colonization induces higher nasal bacterial loads than adult colonization

Next, we compared *S. aureus* colonization patterns between three groups of age-matched mice: (i) neonatally colonized mice, (ii) mice colonized as adults, and (iii) PBS-treated controls. Neonatal colonization was achieved by vertical transmission, while adults were colonized by intranasal inoculation with 10^8^ CFU JSNZ [29,33] at the age of 8 weeks. In the mice colonized as adults, bacterial loads were determined 6, 14, and 28 days post-colonization; animals colonized as neonates were sampled as adults at the same age of 9-13 weeks. Adult colonization resulted in high bacterial loads in the nose, cecum, and feces at days 6 and 14 (Fig 2A-C), consistent with an independent experiment examining loads in adult mice at days 3, 6, and 10 post-colonization (S1 Fig). At day 28, however, 5 of 11 mice in the adult group had cleared *S. aureus* from the nose while remaining colonized in the gut (Fig 2). In contrast, neonatal colonization was stable, with high bacterial loads in nose and gut in almost all animals through day 28 (Fig 2). Thus, natural neonatal exposure to *S. aureus* JSNZ results in long-term stable carriage, whereas experimental adult colonization is less robust.

**Fig 2.**
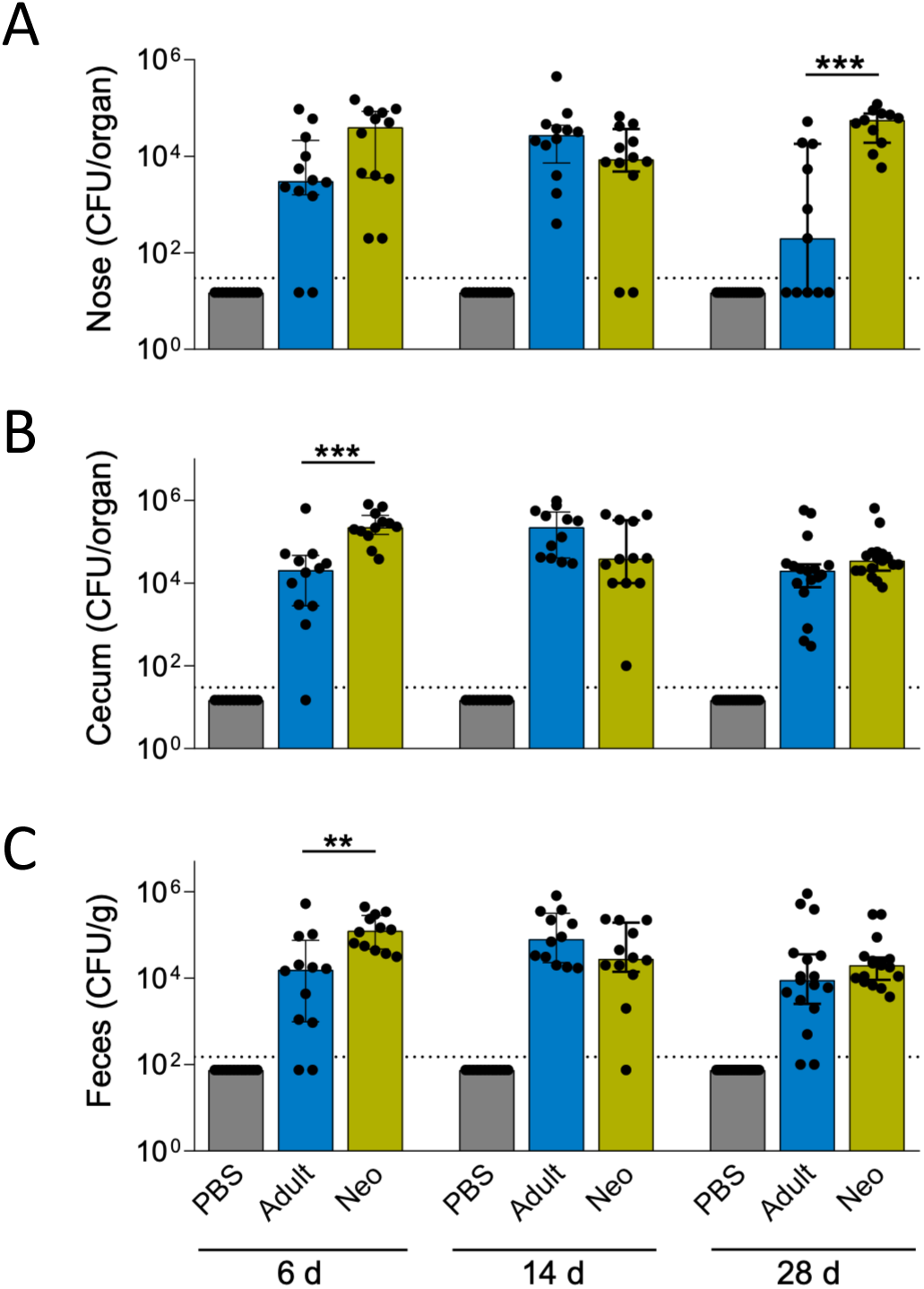
Neonatal exposure to *S. aureus* JSNZ results in more persistent colonization than adult exposure. Female C57BL/6N mice were inoculated i.n. with 1 × 10⁸ CFU *S. aureus* JSNZ at 8 weeks of age (adult colonization, Adult) or colonized naturally by vertical transmission after birth (neonatal colonization, Neo). PBS-inoculated mice served as controls. Bacterial loads in the nose (A), cecum (B), and feces (C) were determined at the indicated time points post-colonization of adult mice by homogenization and plating on selective agar. Dashed line indicates the lower limit of detection), negative samples are depicted as LLOD/2. Data are combined from two independent experiments (n = 11 - 17 mice/group) and presented as median with IQR. Statistics: Mann-Whitney U test. CFU, colony-forming units.

### Neonatal colonization inhibits chemokine production in the nose

Nasal *S. aureus* colonization induces an early local chemokine response that contributes to eliminating nasal carriage [21,39–41]. Consistent with this, we observed an early increase in RANTES, MIP-1α, and KC at day 3 post-colonization in adults compared to PBS-inoculated controls, which returned to baseline by day 6 (S2 Fig). The subsequent experiment comparing adult and neonatal colonization did not include an early time point. As expected, levels were similar in the adult colonization and PBS control groups at day 6 post-colonization. In contrast, neonatally colonized mice exhibited lower levels of chemokines involved in recruiting Th1 cells (RANTES, MIP-1α), Th17 cells (MIP-3α), Th2 cells (MCP-1, Eotaxin, TARC), B cells (BLC), and neutrophils (KC) (Fig 3) [42]. Local cytokine levels did not differ significantly between groups (data not shown). We conclude that reduced chemokine levels may have favored prolonged *S. aureus* persistence following neonatal colonization.

**Fig 3.**
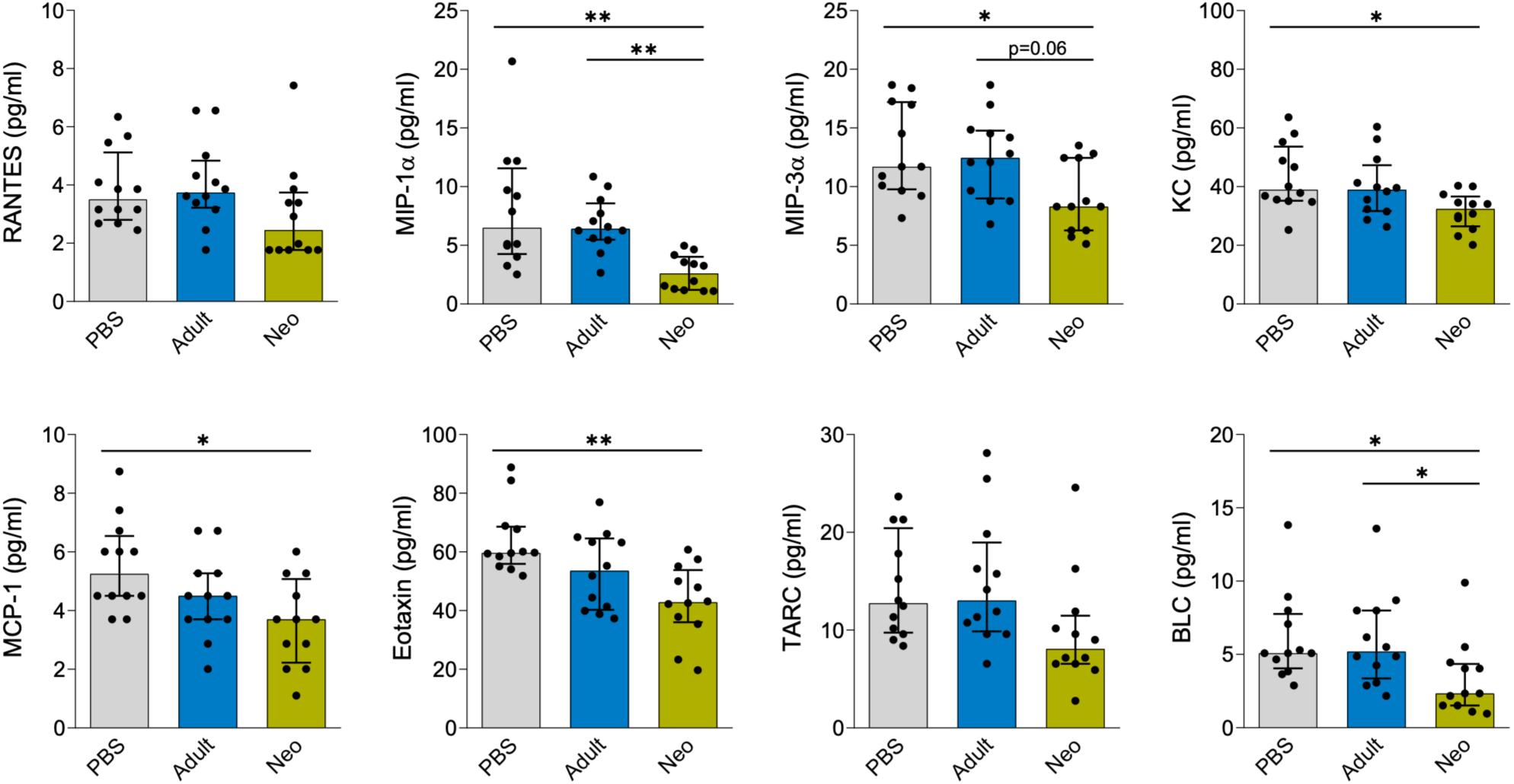
Neonatal *S. aureus* colonization is associated with reduced nasal chemokine levels. Chemokine concentrations in nasal homogenates were measured by bead-based multiplex assay in female C57BL/6N mice colonized with *S. aureus* JSNZ as adults (day 6 post-colonization), age-matched neonatally colonized mice, and PBS-inoculated controls. Chemokines not differing significantly between groups (CXCL10, CCL4, CXCL9, LIX, and MDC) are not shown. Data are presented as median with IQR (n = 12 mice/group). Statistics: Kruskal-Wallis test with Dunn’s multiple comparisons test. Chemokine aliases: RANTES (CCL5), MIP-1α (CCL3), MIP-3α (CCL20), KC (CXCL1), MCP-1 (CCL2), Eotaxin (CCL11), TARC (CCL17), BLC (CXCL13).

### *S. aureus* nasal colonization generates *S. aureus*-specific Th17 memory cells

T cells are important in controlling *S. aureus* nasal colonization in humans [43–45] and in murine models of transient colonization [21–23,32,46]. Here, we characterized the *S. aureus*-specific T cell response to persistent neonatal versus adult colonization and identified the responding T cell subpopulations. CFSE-labelled cervical lymph node (CLN) cells were left unstimulated or re-stimulated *in vitro* with an *S. aureus* antigen cocktail (heat-inactivated extracellular proteins and UV-inactivated JSNZΔ*spa* cells) for 4 days. Antigen-specific T cell activation was assessed using three complementary readouts: (1) flow cytometry-based phenotyping of T cell subsets and proliferation, (2) cytokine quantification in culture supernatants, and (3) enumeration of cytokine-producing cells. The quality of the anti-*S. aureus* T cell response (Th subsets and memory subtypes) was analyzed by flow cytometry (gating strategy in S3 Fig). Re-stimulation with *S. aureus* antigens induced an expansion of Th17 cells (RORγt^+^CD4^+^) in both neonatal and adult colonization groups but not in PBS-inoculated controls. This was reflected by a higher percentage of Th17 cells within the total CD4^+^ T cell population (Fig 4A, C left panel) and among the proliferating effector/effector memory T helper cells (CD4^+^RORγt^+^CD44^+^CD62L^−^CFSE^dim^) (Fig 4B) upon re-stimulation at days 6, 14, and 28. For instance, in the control group only 6.0% of the proliferating effector memory CD4+ T cells were RORγt^+^ upon re-stimulation, as compared to 34.2% in adult and 26.0% in neonatal colonization (Fig 4C, middle panel).

**Fig 4.**
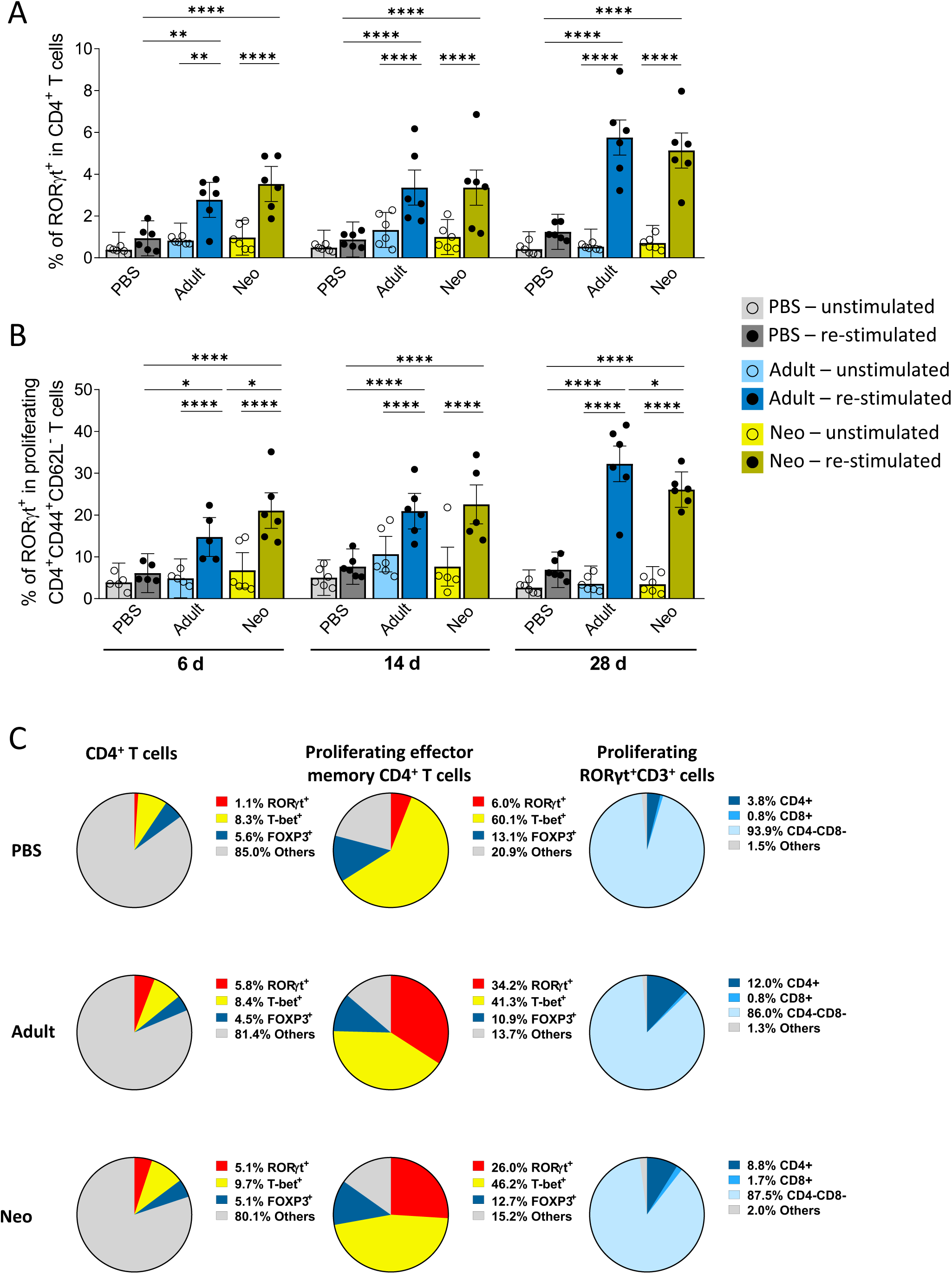
*In vitro* re-stimulation induces *S. aureus*-specific Th17 cell expansion in adult and neonatally colonized mice. CLN cells from adult or neonatally colonized mice and PBS-inoculated controls were harvested at the indicated time points post-colonization of adult mice, labelled with CFSE, and cultured for 4 days in the presence or absence of an *S. aureus* antigen cocktail. T cell subpopulations and proliferation were assessed by flow cytometry. Bar charts show the percentage of RORγt^+^ cells within the total CD4^+^ T cell population (CD3^+^CD4^+^CD8^−^) (A) and within the proliferating effector memory CD4^+^ T cell population (CFSE^low^CD3^+^CD4^+^CD8^−^CD44^+^CD62L^−^) (B). Each dot represents pooled CLN cells from two mice (n = 10 - 12 mice/group). Pie charts show the median proportion of RORγt^+^, T-bet^+^, and FOXP3^+^ cells within the total CD4^+^ T cell population (left) and the proliferating effector memory CD4^+^ T cell population (middle), and the proportion of CD4^+^, CD8^+^, and CD4^−^CD8^−^ T cells among proliferating RORγt^+^ T cells (right; median of 6 samples/group) upon re-stimulation (C). Data in (A) and (B) are presented as mean ± confidence interval (CI). Statistics: linear mixed model (including animal-ID as random factor) in two-way ANOVA design.

Similarly, the proportion of CD4^+^ T cells within the proliferating RORγt^+^CD3^+^ T cell population was higher in adult and neonatal colonization (12.0% and 8.8%) than in control animals (3.8%) (Fig4C, right panel). These data demonstrate that RORγt^+^CD4^+^ T cells were primed in the cervical lymph nodes during nasal colonization and expanded upon re-challenge with *S. aureus* antigens. Notably, this *S. aureus*-specific Th17 memory response was not restricted to the draining lymph nodes but was also detected in the spleen (S5 Fig).

By contrast, for Th1 cells (T-bet+CD4+) we observed only a moderate and transient increase in antigen-specific CD4^+^T cells at 6 days in the adult colonization group (S4 Fig). However, there was no increase in T-bet^+^ cells within proliferating effector/effector memory Th cells at 28 days (Fig4C, middle panel). No clear expansion of Treg (CD4^+^FOXP3^+^) cells was detectable within the total CD4^+^ T cell population and any time point (S4 Fig). In line with this, colonization did not affect the percentage of Tregs among the proliferating effector/effector memory Th cells at 28 days (Fig4C, middle panel).

Collectively, flow cytometry data demonstrate that *S. aureus* colonization triggers a strong, lasting *S. aureus*-specific Th17 response (persisting for at least 28 days), with no appreciable difference between neonatal and adult colonization, when both groups are sampled as adults.

### *In vitro* re-stimulation induces *S. aureus*-specific Th17-, Th1-, Th2-, and Treg-associated cytokine production

To further characterize the cellular immune response to neonatal and adult colonization, cytokine concentrations in cell culture supernatants and the number of cytokine-producing cells were assessed upon *in vitro* re-stimulation with the *S. aureus* antigen cocktail. Re-stimulation induced a strong cytokine response at all time points, with mostly similar patterns in the adult and neonatal groups, while cells from the PBS controls did not respond.

CLN cells (Fig 5) and splenocytes (S6 Fig) from both colonized groups secreted substantially elevated amounts of Th17-associated cytokines (IL-17A, IL-17F, and IL-22) upon re-stimulation at all time points compared to PBS controls. This pattern was also observed at the early stage of adult colonization (day 3; S7 Fig). CLN cells from adult and neonatally colonized mice produced relatively high levels of IFN-γ spontaneously, which were elevated upon re-stimulation at all time points (Fig 5). Th2-associated cytokines (IL-5 and IL-13) were also considerably increased in re-stimulated CLN cells and splenocytes from both colonized groups as compared to the PBS controls. Notably, CLN cells and splenocytes from neonatally-colonized mice released higher IL-5 levels than adult-colonized mice at all time points. The regulatory cytokine IL-10 was also released by re-stimulated T cells from both colonization groups (Fig 5, S6 Fig). Elevated IL-10 levels were also observed at the early stage of adult colonization (day 3; S7 Fig).

**Fig 5.**
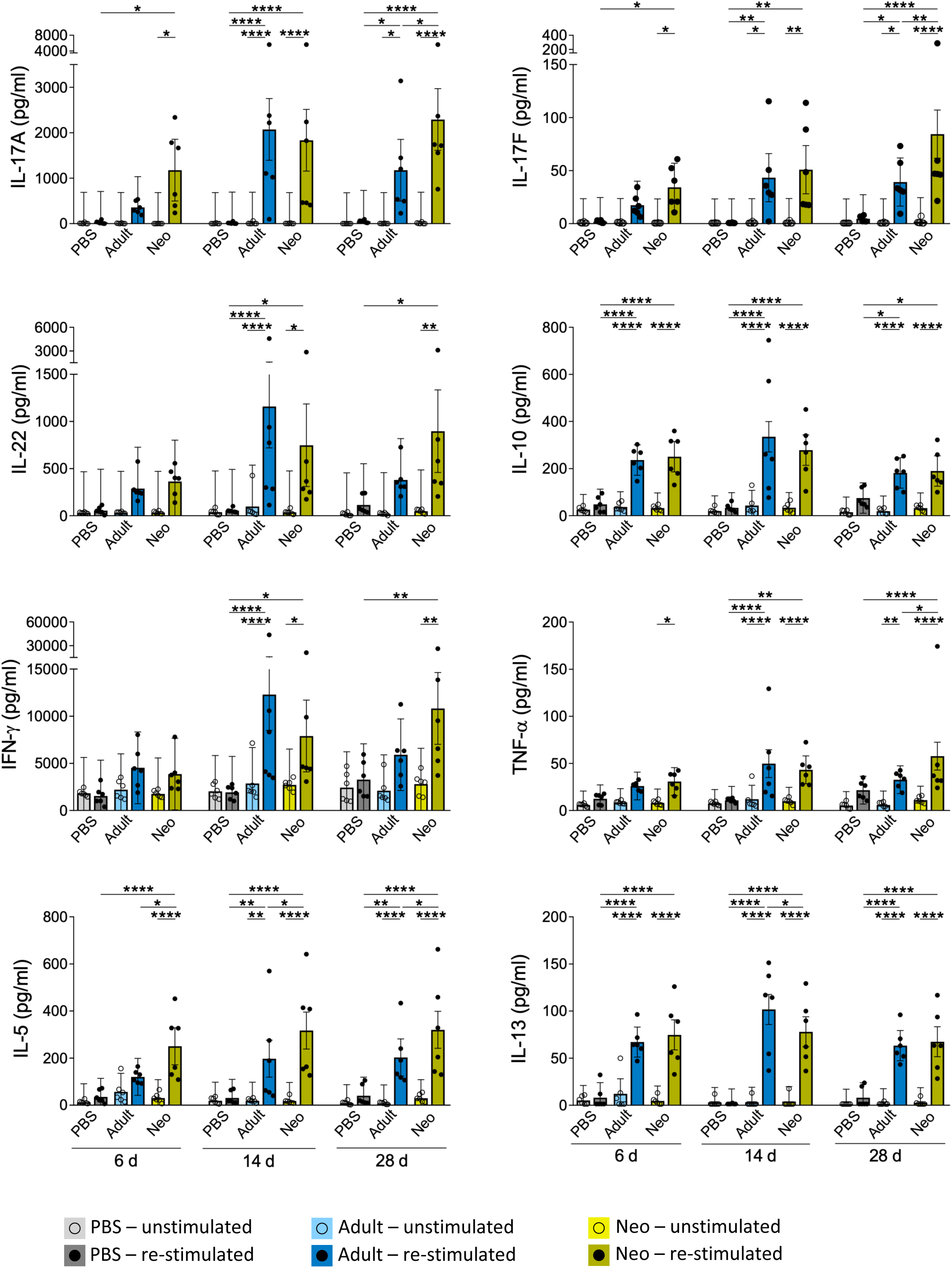
*S. aureus-*specific CLN cells from colonized mice secrete high amounts of IL-17A. CLN cells from adult-or neonatally colonized mice and PBS-inoculated controls were cultured for 4 days in the presence or absence of an *S. aureus* antigen cocktail. Cytokine concentrations were measured by bead-based multiplex assay or IFN-γ ELISA. Each dot represents pooled CLN cells from two mice (n = 12 mice/group). Data are presented as mean ± CI. Statistics: linear mixed model (including animal-ID as random factor) in two-way ANOVA design.

Next, we enumerated IL-17-, IFN-γ-and IL-10-producing cells upon *in vitro* re-stimulation using a FluoroSpot assay. The number of IL-17A-producing cells was strongly increased upon re-stimulation in both colonized groups compared to controls (Fig 6A). Consistent with the cytokine measurements (Fig5), there were high numbers of spontaneously IFN-γ-secreting cells in all groups. These increased moderately in the colonized groups upon re-stimulation at days 6 and 14, the difference reaching significance in both groups at day 28 (Fig 6B). Re-stimulation similarly increased IL-10-secreting cell numbers after both neonatal and adult colonization.

**Fig 6.**
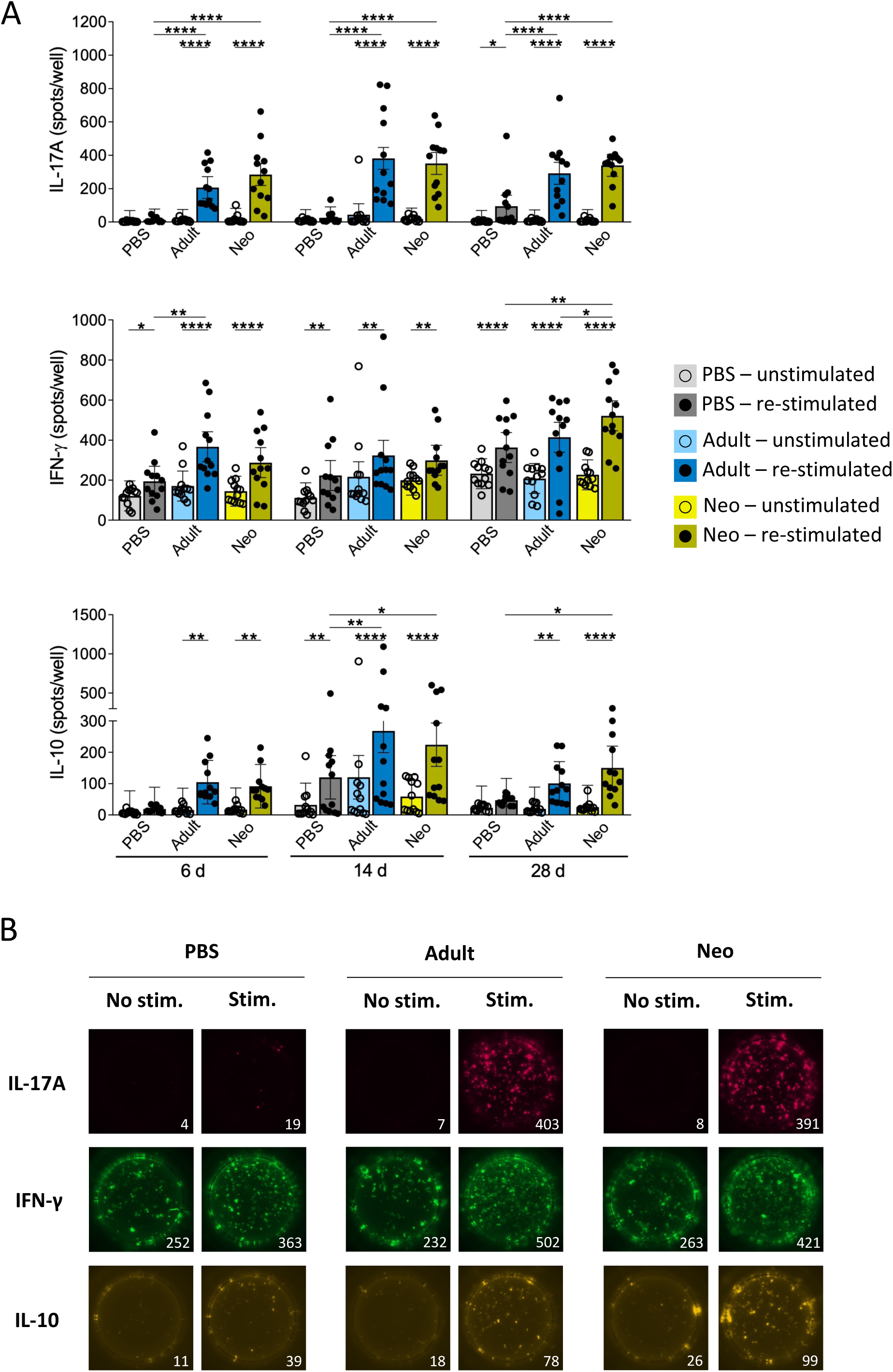
*S. aureus* colonization induces IL-17A-and IL-10-secreting CLN cells. CLN cells from adult-or neonatally colonized mice and PBS-inoculated controls were cultured for 3 days in pre-coated FluoroSpot plates in the presence or absence of an *S. aureus* antigen cocktail. IL-17A-, IFN-γ-, and IL-10-secreting cells were enumerated using a FluoroSpot reader (A). Representative images from day 28 of colonization are shown (B). Data are combined from two independent experiments (n = 12 mice/group) and presented as mean ± CI. Statistics: linear mixed model (including animal-ID as random factor) in two-way ANOVA design.

In summary, *in vitro* re-stimulation of CLN cells with the *S. aureus* antigen cocktail induced a Th17-dominated antigen-specific T cell response in JSNZ-colonized animals. Although flow cytometry showed no clear expansion of Th1 or Treg cells, enhanced secretion of type 1, 2, and regulatory cytokines was detected both in culture supernatants and by FluoroSpot.

### IL-17A is mainly produced by Th17 cells

To identify which *S. aureus*-specific T cell subset produced IL-17A upon *in vitro* re-stimulation, CD4^+^, CD8^+^ and γδ T cells were isolated from CLN of colonized mice at 28 days and co-cultured with DCs from *S. aureus*-free mice in the presence or absence of the *S. aureus* antigen cocktail. The purity of isolated CD4^+^ and CD8^+^ T cells was higher than 90%, while the purity of isolated γδ T cells was around 70% (S8 Fig).

Consistent with our previous experiments, CLN cells from both adult and neonatally colonized mice produced high levels of IL-17A upon re-stimulation (Fig 7A). Cell fractionation revealed that IL-17A was predominantly produced by CD4^+^ αβ T cells, while secretion by CD8^+^, and γδ T cells was negligible (Fig 7B).

**Fig 7.**
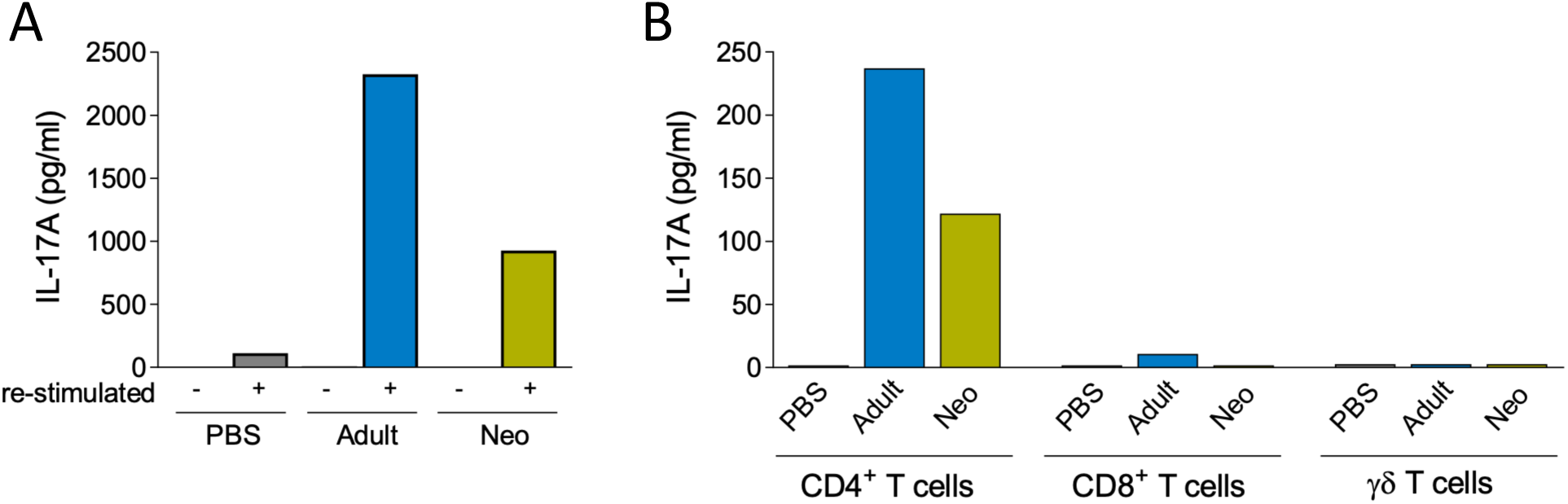
IL-17A production at day 28 post-colonization is predominantly attributed to CD4^+^ T cells. CLN cells were harvested at day 28 from adult-or neonatally colonized mice and PBS-inoculated controls. CLN cells were pooled (5 mice/pool), and CD4^+^, CD8^+^, and γδ T cells were isolated by magnetic bead separation. Unfractionated CLN cells (1 × 10^6^ cells/well) were cultured in the presence or absence of an *S. aureus* antigen cocktail (A). Isolated CD4^+^, CD8^+^ (both 4 × 10⁵ cells/well), and γδ^+^ (1 × 10⁵ cells/well) T cells were co-cultured with dendritic cells (DCs) from naïve female C57BL/6N mice at a DC:T cell ratio of 1:10 and stimulated accordingly (B). Cytokine concentrations were measured by bead-based multiplex assay.

## Discussion

*S. aureus* colonization plays a key role in the epidemiology and pathogenesis of *S. aureus* infections, because most infections have an endogenous origin [4,47,48]. A better understanding of the immune processes involved in colonization could identify protective mechanisms and facilitate vaccine development. This requires animal models that mimic natural and persistent *S. aureus* carriage. Here, we have established a neonatal persistent *S. aureus* colonization model involving the vertical transmission of the mouse-adapted *S. aureus* strain JSNZ from colonized breeding pairs to their offspring. We observed higher bacterial loads and prolonged nasal persistence of JSNZ in neonatal versus adult colonization that was associated with reduced chemokine levels in the nasal tissue. In both groups, colonization elicited a strong Th17-skewed T cell response against the colonizing *S. aureus* strain.

### Vertical transmission of *S. aureus* JSNZ from colonized breeding pairs induces natural, persistent neonatal colonization

In this study, we used the mouse-adapted JSNZ strain to establish an *S. aureus*-positive C57BL/6N breeding colony, obtaining naturally colonized offspring (neonatal colonization). This approach was based on our previous observation that parental animals transferred their colonizing *S. aureus* strain very efficiently to their offspring, resulting in bacterial persistence for months [31]. Maternal transfer models have also been reported for *E. coli* and group B streptococci [49,50].

All breeding animals remained colonized in the nose and gastrointestinal tract throughout the observation period (18-47 weeks). Similarly, human nasal carriage is associated with intestinal *S. aureus* carriage [10]. Notably, the offspring were colonized with high bacterial loads in the nose and gastrointestinal tract for up to 69 weeks, reflecting lifelong natural persistent carriage. Consistent with our findings, Flaxman et al. observed persistent gastrointestinal colonization (up to 3 months) following environmental exposure of neonatal or adult mice to a mouse-adapted *S. aureus* CC15 strain [51,52]. Neonatal colonization models require no manipulation of newborns and are straightforward to implement, eliminating the need for high-dose intranasal inoculation that may mask key aspects of the host immune response in the early phase of natural colonization.

Our neonatal colonization model closely resembles *S. aureus* persistence in human neonates. Human colonization occurs in the first weeks of life, with the mother as the most common source [12,13]. High neonatal susceptibility reflects limited bacterial competition by the developing microbiome [10], which probably contains fewer *S. aureus* inhibiting species such as *Bacillus subtilis* [18], *S. epidermis* [53], *S. lugdunensis* [54] and *Streptococcus spp* [55]. A second facilitating factor is the tolerogenic immune phenotype of neonates [16,17,56]. While approximately 25% of *S. aureus*-positive human neonates clear the bacteria within the first 14 months of life [12], colonization persists beyond one mouse year in our model, likely due to the limited microbial diversity in laboratory mice and continuous intra-cage re-exposure, mimicking human household transmission [57]. Thus, as in in the human situation, the model does not discriminate between persistence and repeated re-colonization from exogenous or endogenous S. aureus reservoirs. But clearly it provides a robust platform for interrogating early-life acquisition of and long-term colonization by *S. aureus* as well as the immune response to the bacteria. Moreover, it may illuminate factors distinguishing *S. aureus* carriers from non-carriers and the impact of colonization on later infections.

### Higher bacterial loads and colonization rate in neonatal vs. adult colonization

In the side-by-side comparison, the adult colonization model was less robust than the neonatal model. Some JSNZ-colonized adult mice eliminated *S. aureus* from the nose by day 28 but remained colonized in the gastrointestinal tract, while neonatal colonization resulted in persistent, high bacterial loads in both niches. This confirms and extends previous studies showing JSNZ to be an excellent short-term (<10 days) colonizer in adult C57BL/6 mice [29,33], exceeding most human-adapted strains in nasal persistence [39]. In contrast, Sun et al. reported 100% colonization with JSNZ in adult BALB/c mice for up to 42 days — differences likely attributable to mouse strain-dependent variation in innate and adaptive immunity or the microbiome [58–61].

### Reduced chemokine levels favor persistent neonatal colonization

To assess whether differential immune cell recruitment contributed to the distinct colonization patterns, we investigated local chemokine levels. In line with other studies [21,39], adult colonization elicited an early increase (day 3) in RANTES, MIP-1α, and KC compared to PBS controls, returning to baseline by day 6. In contrast, neonatal colonization reduced nasal chemokine levels below baseline on day 6. Since chemokines are pivotal for immune cell recruitment, reduced levels likely contributed to long-term *S. aureus* persistence. Ge et al. similarly showed that colonization duration correlates with the early local chemokine response and neutrophil recruitment: the short-term colonizer Newman upregulated nasal pro-inflammatory chemokines at 3 hours post-colonization and triggered neutrophil influx, whereas the persistent strain JKD6159 did not [39]. Corroborating this, neonatal macrophages show reduced chemokine production including KC and MCP-1 upon stimulation with killed pneumococci [62], and circulating neutrophils from infant mice express lower levels of the chemokine receptor CXCR2 than those from adults [63]. Collectively, reduced local chemokine levels likely impaired neutrophil recruitment to the nasal compartment, promoting colonization and persistent *S. aureus* carriage in neonates.

### Colonization induces S. aureus-specific T cell response, dominated by Th17 cells

While previous studies used knock-out mouse strains to study T cell roles in *S. aureus* colonization, we employed an *S. aureus* antigen cocktail to decipher antigen-specific T cell responses. The JSNZ strain provides a broad antigen spectrum but lacks superantigens [29,31,64], which would otherwise interfere with this analysis. This approach therefore recalls immune responses against multiple *S. aureus* antigens but may fail to fully reproduce dynamic antigen expression, immunomodulatory activity of virulence factors, or local environmental conditions during colonization *in vivo*. These limitations may partly explain why no major differences in cellular immune phenotypes or cytokine profiles were observed between the groups colonized as neonates or adults.

Overall, we observed a Th17-dominated T cell response alongside cytokine profiles consistent with the response of Th1, Th2, and Treg cells. In JSNZ-colonized male mice that spontaneously developed a purulent infection of their preputial glands we also found Th1-, Th2-, and Treg-associated cytokines along with an even more pronounced Th17 cytokine profile [38]. These findings are in line with a previous study showing strong antigen-specific proliferation and robust IL-17A production, with lower IFN-γ and IL-5 production, upon re-stimulation of lymph node cells from *S. aureus*-exposed mice [65].

Using multiple read-outs, we demonstrated that persistent nasal colonization primes a Th17-dominated response. *S. aureus*-specific RORγt⁺ Th17 memory cells in draining lymph nodes responded to *in vitro* re-stimulation with proliferation and the release of large amounts of IL-17A, with CD4⁺ T helper cells as the main IL-17A source in both groups. Notably, *S. aureus*-reactive Th17 cells were also detected in the spleen, suggesting subclinical systemic spread of the bacteria and/or migration of the immune cells beyond the nasal and gut mucosa.

Th17 cells are protective in *S. aureus* colonization and infection [21–23,44,46]. IL-17 promotes decolonization via antimicrobial peptide expression and neutrophil influx [21,22,40]. Similarly, *S. aureus* exposure of human nasopharynx-associated lymphoid tissues induces Th17 responses [66]. IL-17-producing γδ T cells also mediate protection [23,67–69], acting as a major IL-17 source at days 3 and 7 post-colonization in mice [23]. However, in a gastrointestinal colonization model Th17 cells and neutrophils — not γδ T cells — drove clearance [61]. Nonetheless, this IL-17A-dominant recall response was insufficient for *S. aureus* clearance in our model. This is in agreement with human data linking a high Th17:Th1 mRNA ratio to persistent nasal colonization [44,70].

High levels of IFN-γ and TNF-α were also measured after re-stimulation, but there was no expansion of T-bet⁺ Th cells, suggesting T-bet-independent Th1 cells [71,72] and CD8⁺ T cells as likely IFN-γ sources. Memory CD8⁺ T cell responses were described in *S. aureus*-exposed humans and mice [70,73]. While Th1 responses contribute to clearance of gastrointestinal colonization [46], their role in nasal colonization remains incompletely understood.

Immune cells of the CLNs further elaborated type 2 cytokines upon re-stimulation with *S. aureus* antigens, mirroring findings in healthy humans who harbor *S. aureus*-specific memory T cells that secrete Th2 cytokines [24,73,74]. *S. aureus* can promote Th2 skewing via secreted allergens, proteases and superantigens. This is considered as an immune evasion strategy [24,74,75] because of the association between atopic disorders and *S. aureus* colonization [75,76]. Remarkably, IL-5 was the only cytokine whose secretion was significantly higher in neonatally than in adult-colonized mice. In our colonization models, we did not observe Foxp3⁺ T cell expansion, but there was pronounced IL-10 secretion by lymph node cells and splenocytes after re-stimulation. Notably, IL-10 was already elevated upon re-stimulation at day 3 post-colonization, suggesting innate memory reactivation. IL-10 plays a dual role in the *S. aureus*-host balance: suppressing effective defense during colonization and biofilm formation and limiting harmful inflammation in acute systemic infection [23,77–79]. IL-10 can be produced by Tregs, macrophages, MDSCs, and Bregs [25,80]. Kelly et al. reported IL-10 upregulation in myeloid cells and B cells in the nasal cavity of mice 3 -7 days post-colonization, which facilitated persistence of *S. aureus* by suppressing local IL-17 and IL-22 responses [23]. These anti-inflammatory responses to *S. aureus* may be driven by bacterial PAMPs and context-dependent virulence factor regulation. They probably influence the outcome of infection and vaccination, warranting further investigation [79,81–83].

### Persistence mechanisms in neonatal vs. adult colonization

Given that mice colonized with *S. aureus* as neonates were persistently colonized and that their nasal bacterial loads were 10-100fold higher than those in animals colonized as adults, we speculated that the tolerogenic immune status of newborns might underlie this difference. Neonatal exposure to commensal microbes and pathobiont bacteria can induce features of immune tolerance [16,56,56,84–87], and in a murine MRSA pneumonia model neonatal mice had worse survival, less inflammation and reduced T cell responses compared to adults [88]. However, we found no significant differences in regulatory T cell responses between the JSNZ-colonized groups - neither in Treg subtype expansion nor in cytokine secretion. In contrast, we observed a moderately stronger Th2 response in the neonatally colonized mice, which was significant in the case of IL-5. This supports the notion that early in life exposure to *S. aureus* induces a Th2-polarization that may contribute to bacterial persistence by counterbalancing Th1/17-driven inflammation. Moreover, animals that were colonized as neonates had significantly reduced chemokine levels locally in the nose when tested as adults.

It is possible that we missed a subtle Treg-phenotype in our model of neonatal colonization. First, the complex antigen cocktail used for in vitro re-stimulation may have activated non-T cells, masking neonatal immunosuppressive features. Second, rare antigen-specific T cell subsets may have escaped flow cytometric detection. Third, beyond Tregs, MDSCs play key roles in *S. aureus* infections [78,89] but were not addressed here. Finally, early *S. aureus* exposure may shape microbiome composition and thereby indirectly modulate immune activity. In the future, studies of the microbiome combined with single-cell or OMICs approaches [90] will complement our findings and advance understanding of the memory T cell response generated during *S. aureus* colonization.

### Summary

Our neonatal colonization model in mice is well-suited for dissecting host-pathogen interactions during natural persistent *S. aureus* carriage. Its utility extends beyond this article - from improving the translational value of preclinical vaccine and therapeutic studies, to advancing our understanding of host-*S. aureus* interactions, including virulence factor regulation during colonization, the shift to pathogenicity, and host and bacterial determinants of carriage. Application should be guided by the specific research question and interpreted in light of the model’s inherent limitations, particularly regarding species-specific targets of virulence factors.

This work also provides important insights into the adaptive immune response to *S. aureus* colonization. We demonstrate for the first time in mice that nasal colonization induces a strong, long-lasting Th17-dominated memory response poised for rapid secondary recall. The protective capacity of these memory Th17 cells is likely tempered *in vivo* by IL-10-producing regulatory cells and Th2-biased priming. Since humans are naturally colonized by *S. aureus*, such mechanisms will influence their outcomes of infection and vaccination and warrant future investigation.

## Material and Methods

### *S. aureus* strains and growth conditions

Murine *S. aureus* colonization experiments used the mouse-adapted *S. aureus* strain JSNZ (ST88/CC88-MSSA), isolated from a C57BL/6J mouse colony [29]. This strain lacks superantigens, exfoliative toxins, Panton-Valentine leucocidin (PVL) and immune evasion cluster (IEC) genes. JSNZ was grown in Brain Heart Infusion medium to the mid-logarithmic phase and prepared for intranasal inoculation as previously described [33]. A protein A-deficient JSNZ mutant (JSNZΔ*spa*) was used for *in vitro* re-stimulation assays [31].

### *S. aureus*-free and -colonized C57BL/6NRj colonies

Male and female C57BL/6NRj mice with Specific and Opportunistic Pathogen Free status (SOPF, *S. aureus*-free, 6-8 weeks old) were purchased from Janvier Labs (Saint-Berhevin, France) and mated to establish a *S. aureus*-free breeding colony. Stool samples were collected regularly from every cage and screened for *S. aureus*.

To obtain naturally *S. aureus*-colonized mice, male and female C57BL/6NRj mice (7 weeks old) were intranasally (*i.n.*) inoculated with 1 × 10^8^ CFU *S. aureus* JSNZ under mild anesthesia (2 L/min O_2_ and 2% isoflurane). One week post colonization, mice were mated and the offspring were weaned at 21 days. *S. aureus* colonization of breeding pairs and offspring was monitored regularly by fecal load determination. The *S. aureus*-colonized colony was maintained in a separate S2-classified animal facility.

Both colonies were housed under identical SOPF conditions in Individually Ventilated Cages (IVC, 4 animals/cage) with autoclaved bedding, enrichment, and nesting materials. Autoclaved food and water were provided *ad libitum*.

### Adult and neonatal *S. aureus* colonization models

For the adult colonization, *S. aureus*-free female C57BL/6NRj mice (8 weeks old) were colonized *i.n.* with 1 × 10^8^ CFU *S. aureus* JSNZ in 10 µl (5 µl/nostril) under mild anesthesia (2% isoflurane and 2 L/min O_2_). For neonatal *S. aureus* colonization, female mice were naturally colonized via vertical transmission from JSNZ-positive breeding pairs. For comparison with mice colonized as adults, age-matched neonatally colonized mice were used. Age-matched PBS-inoculated mice (5 µl/nostril) served as controls. Experiments were performed in replicates using 4-6 animals/group. Fecal samples were collected individually before colonization and on the day of read-out (day 6, 14 and 28 post colonization of the adult group). In a separate experiment, adult colonization (1 × 10^8^ CFU *S. aureus* JSNZ) was compared to PBS-inoculated controls on days 3, 6, and 10 post-colonization.

The animals were anesthetized at the indicated days with ketamine/xylazine (100 mg/10 mg per Kg body weight), and blood samples were collected via the retro-orbital plexus, followed by terminal isoflurane overdose. Nasal tissue (nasoturbinate, maxilloturbinate and parts of the ethmoid turbinate(s)) and cecum were harvested and stored at −80°C for bacterial load quantification. Spleen and cervical lymph nodes (CLN) were collected in 5% fetal bovine serum (FBS) (Sigma-Aldrich, St. Louis, USA)/phosphate-buffered saline (PBS, Pan Biotech, Aidenbach, Germany) for the analysis of *S. aureus*-specific T cells.

### Determination of the bacterial load

Bacterial load in nose, cecum and feces was determined as previously described [33]. Briefly, samples were homogenized in PBS containing protease inhibitor cocktail (cOmplete tablets, Mini EDTA-free, EASYpack, Roche, Basel, Switzerland) and 10 µl of serial dilutions were plated in triplicate on *S. aureus* CHROMagar plates (CHROMagar, Paris, France). After incubation at 37°C for 24 h, colony-forming units (CFUs) were counted.

### Preparation of single cell suspensions from tissues

Spleens and CLN were passed through a 70 µm cell strainer, washed with 25 ml 5% FBS/PBS, and centrifuged at 300 × *g* for 8 min at 4°C. CLN cells were resuspended in 400 µl of 5% FBS/PBS. Splenocytes were resuspended in 2 ml RBC lysis buffer (155 mM NH₄Cl, 10 mM KHCO₃, 0.1 mM EDTA in distilled water; pH 7.3) for 2 min at room temperature (RT), then quenched with 40 ml of 5% FBS/PBS and centrifuged at 300 × *g* for 8 min at 4°C. After discarding the supernatant, splenocytes were resuspended in 3 ml of 5% FBS/PBS. Cell concentration was determined using Trucount™ beads (BD Biosciences, San Diego, USA).

### Magnetic bead isolation of T cell subsets

Gamma-delta (γδ)^+^, CD4^+^, and CD8^+^ T cells were isolated from CLN cell suspensions by a two-step isolation magnetic bead procedure. In the first step, TCRγ/δ^+^ T cells were isolated using a mouse TCRγ/δ^+^ T cell isolation kit (Miltenyi Biotec, Bergisch Gladbach, Germany). In the second step, CD4^+^ and CD8^+^ T cells were enriched from the total TCRγ/δ^−^ cell fraction by positive and negative selection, respectively, using CD4 (L3T4) Microbeads (Miltenyi Biotec).

### CFSE labelling

To assess cell proliferation, splenocytes and CLN cells were labelled with carboxyfluorescein diacetate succinimidyl ester (CFDA-SE; BioLegend, San Diego, USA) according to the manufacturer’s protocol. Briefly, 4 × 10⁶ cells in 1 ml PBS were incubated with CFDA-SE at a final concentration of 4 µM for 5 min at 37°C in the dark. Cells were washed with 40 ml of 10% FBS/PBS by centrifugation at 300 × *g* for 8 min at RT, followed by a second wash in 10 ml TexMACS medium (Miltenyi Biotec) supplemented with 10% FBS, 1 × Penicillin/Streptomycin/Glutamine, and 50 µM β-mercaptoethanol (T10F medium). CFSE-labelled cells were resuspended in T10F medium at 1 × 10⁷ cells/ml for subsequent culture experiments.

### Cell culture

CFSE-labelled cells were seeded in duplicate at 1 × 10⁶ cells/well. Isolated CD4^+^, CD8^+^ (both 4 × 10⁵) and γδ^+^ T cells (1 × 10⁵) were co-cultured with BMDCs at a DC:T cell ratio of 1:10. Unfractionated cells were seeded at 1 × 10⁶/well. All cells were cultured in 200 µl T10F medium supplemented with 20 ng/ml IL-2 (BioLegend) in 96-well round-bottom microplates at 37°C and 5% CO₂ for 4 days, either without stimulation or stimulated with an *S. aureus* antigen cocktail consisting of heat-inactivated extracellular proteins (5 µg/ml) and UV-inactivated cells of strain JSNZΔ*spa* (2.5 × 10⁵ CFU/ml). Afterwards, cells were harvested by centrifugation at 300 × *g* for 8 min at 4°C and subjected to flow cytometry analysis; cell-free supernatants were stored at −80°C for cytokine measurements.

### Flow cytometry

CFSE-labelled cells were washed twice with 2 ml PBS by centrifugation at 300 × *g* for 8 min at 4°C, then incubated with Zombie NIR™ (BioLegend) for 30 min at RT in the dark. After washing with 2 ml FACS buffer (PBS containing 2% FBS, 0.02% sodium azide, and 2 mM EDTA), cells were incubated with FcR blocking solution (Miltenyi Biotec) for 5 min at 4°C. Cells were then surface-stained for 20 min at 4°C with antibodies against CD3 (BV650), CD4 (BV605), CD8 (BV510), CD62L (PerCP-Cy5.5), and CD44 (PE-Cy7) (all BioLegend). For intranuclear transcription factor staining, cells were washed, permeabilized, and fixed using the MACS Transcription Factor Buffer Set (Miltenyi Biotec) per the manufacturer’s protocol, followed by intracellular staining for 30 min at 4°C with antibodies against RORγt (PE; BD Biosciences), FoxP3 (VioR667; Miltenyi Biotec), and T-bet (BV421; BioLegend). Gates for all transcription factors were set using Fluorescence Minus One (FMO) controls.

To verify the purity of isolated T cell (CD4^+^, CD8^+^, and γδ^+^) populations, cells were washed with PBS, stained with Zombie NIR™, and blocked with FcR blocking solution as above, then stained for 20 min at 4°C with antibodies against CD3 (PerCP-Vio700), CD4 (VioR720), CD8 (VioGreen), and TCRγδ (PE-Vio770) (all Miltenyi Biotec).

All cells were resuspended in 200 µl FACS buffer containing BD Trucount™ beads. Data were acquired on an LSR II flow cytometer (BD Biosciences) and analyzed using FlowJo™ v10 (BD Biosciences). Antibody details are provided in S1 and S2 Tables.

### FluoroSpot

IFN-γ, IL-17A, and IL-10 secreting cells were quantified by FluoroSpot using mouse detection kits, antibodies, and plates from Mabtech (Nacka, Sweden) according to the manufacturer’s instructions. Briefly, 1 × 10⁶ CLN cells were seeded into pre-coated FluoroSpot plates in 200 µl T10F medium supplemented with 20 ng/ml IL-2 and 0.2 µg/ml anti-CD28 monoclonal antibody, and either left unstimulated or stimulated with the *S. aureus* antigen cocktail described above for 3 days at 37°C and 5% CO₂. Plates were then washed five times with PBS and incubated with the respective detection antibodies and fluorophore-conjugated reagents provided by the manufacturer. Spots were read and counted by Mabtech Consulting using a FluoroSpot reader (Mabtech IRIS™) with Mabtech Apex™ software (version 1.1.9).

### Detection of mouse chemokines and cytokines

Chemokine and cytokine levels in nose homogenates and cell-free culture supernatants were measured using the LEGENDplex™ Mouse Proinflammatory Chemokine Panel (13-plex, BioLegend) and the LEGENDplex™ MU Th Cytokine Panel (12-plex, BioLegend), respectively. The chemokine panel simultaneously quantifies MCP-1 (CCL2), RANTES (CCL5), IP-10 (CXCL10), Eotaxin (CCL11), TARC (CCL17), MIP-1α (CCL3), MIP-1β (CCL4), MIG (CXCL9), MIP-3α (CCL20), LIX (CXCL5), KC (CXCL1), BLC (CXCL13), and MDC (CCL22); the cytokine panel simultaneously quantifies IL-2, −4, −5, −6, −9, −10, −13, - 17A, −17F, −22, IFN-γ, and TNF-α. Both panels were analyzed according to the manufacturer’s instructions on an LSR II flow cytometer (BD Biosciences).

Samples with saturated IFN-γ levels were diluted and reanalysed using the MAX™ Deluxe Set Mouse IFN-γ ELISA kit (BioLegend) according to the manufacturer’s instructions. Absorbance was measured at OD 450 nm using an Infinite M200 Pro microplate reader (Tecan, Crailsheim, Germany).

### Ethics statement

All animal experiments were approved by the local State authorities (Landesamt für Landwirtschaft, Lebensmittelsicherheit und Fischerei Mecklenburg-Vorpommern, 7221.3-1-018/19 and 7221.3-1-058/20). All experiments were performed in accordance with the German Animal Welfare Act (Deutsches Tierschutzgesetz), the EU Directive 2010/63/EU for animal experiments and the Federation of Laboratory Animal Science Associations (FELASA) and comply with the ARRIVE guidelines.

### Statistics

Data analysis was performed using the GraphPad Prism 8 package (GraphPad Software, Inc., La Jolla, USA) and R (core v.4.5.0) [91] for statistical computing (tidyverse package v.2.0.0 [92]; rstatix package v.0.7.2 [93]) and data visualization (ggplot2 package v3.5.2.) [94]. Two groups (skewed distribution) were compared using the Mann-Whitney test. More than two groups were analyzed using a Kruskal-Wallis test with Dunńs test for multiple comparisons (skewed distribution). Data from re-stimulation assays present a combination of paired and unpaired samples and were analyzed by a linear mixed model in two-way ANOVA design.

*P < 0.05, **P < 0.01, ***P < 0.001, ****P < 0.0001. Correlation analyses were performed using Spearman nonparametric correlation and the correlation coefficient r and the p value were reported. Details of statistical analyses, including statistical tests and significance criteria, are provided in the figure legends.

## Acknowledgements

The authors would like to thank Sabine Berg for her support in planning the animal experiments. The authors thank Jens van den Brandt and the team of the ZSFV for their support in animal husbandry. The authors would like to thank Sabine Prettin, Fawaz Alsholui, Susanne Neumeister, and Aneta Martiniuc for technical support.

Portions of this manuscript were edited for language and clarity using Claude (Anthropic, San Francisco, CA, USA). All AI-generated suggestions were reviewed and approved by the authors, who take full responsibility for the content.

---

**S1 Fig. Intranasal inoculation of adult mice with *S. aureus* JSNZ induces stable colonization over 10 days.** Female 8-week-old C57BL/6N mice were inoculated i.n. with PBS or 1 × 10⁸ CFU *S. aureus* JSNZ. Bacterial loads in the nose, cecum, and feces were determined at days 3, 6, and 10 post-colonization by homogenization and plating on selective agar. Dashed line indicates the lower limit of detection), negative samples are depicted as LLOD/2. Data are presented as median with IQR (n = 8-14 mice/group). CFU, colony-forming units.

**S2 Fig. *S. aureus* colonization of adult mice induces an early transient increase in nasal chemokine levels.** Female 8-week-old C57BL/6N mice were inoculated i.n. with PBS or 1 × 10⁸ CFU *S. aureus* JSNZ. Chemokine concentrations in nose tissue homogenates were measured at the indicated time points post-colonization by bead-based multiplex assay. Data are presented as median with IQR (n = 8-12 mice/group). Statistics: Mann-Whitney U test. Chemokine aliases: RANTES (CCL5), MIP-1α (CCL3), MIP-3α (CCL20), KC (CXCL1), MCP-1 (CCL2), Eotaxin (CCL11), TARC (CCL17), BLC (CXCL13), MIP-1β (CCL4), MIG (CXCL9), LIX (CXCL5, and MDC (CCL22).

**S3 Fig. Flow cytometry gating strategy for analysis of the *S. aureus*-specific T cell response after *in vitro* re-stimulation.** CFSE-labelled CLN or spleen single-cell suspensions were cultured for 4 days in the presence or absence of an *S. aureus* antigen cocktail, then stained for surface and intracellular markers. Debris was excluded by FSC/SSC gating, followed by exclusion of Zombie NIR™-positive cells to select live singlets. Live single cells were gated on CD3^+^ to identify T cells, and proliferating T cells were identified as CFSElowCD3^+^. Proliferating RORγt^+^T-bet^−^ cells were gated within the CD3^+^ population. CD4^+^, CD8^+^, and CD4^−^CD8^−^ subsets were gated within total CD3^+^, proliferating CD3^+^, or proliferating RORγt^+^T-bet^−^ T cells. Proliferating cells within each subset were identified by CFSE^low^ gating. Memory subpopulations were defined using CD44 and CD62L: central memory (Tcm, CD44^+^CD62L^+^) and effector memory (Tem, CD44^+^CD62L^−^). Expression of T-bet, RORγt, and FOXP3 was assessed within each T cell subset, proliferating T cells, and proliferating memory T cells. Representative data shown are from re-stimulated CLN cells of the adult colonization group at day 28 post-colonization.

**S4 Fig. *In vitro* re-stimulation of CLN cells does not induce *S. aureus*-specific Th1 or Treg cell expansion following adult or neonatal colonization.** CLN cells from adult-or neonatally colonized mice and PBS-inoculated controls were harvested at the indicated time points, pooled from 2 mice per sample, and cultured for 4 days in the presence or absence of an *S. aureus* antigen cocktail. T cell subpopulations were assessed by flow cytometry. Bar charts show the percentage of T-bet^+^ (Th1) or FOXP3^+^ (Treg) cells within the total Th cell population (CD3^+^CD4^+^CD8^−^). Each dot represents pooled CLN cells from two mice (n = 12 mice/group). Data are presented as mean ± CI. Statistics: linear mixed model (including animal-ID as random factor) in two-way ANOVA design.

**S5 Fig. *In vitro* re-stimulation of splenocytes induces *S. aureus*-specific Th17 cell expansion in adult and neonatal colonization.** Spleens were harvested at the indicated time points from adult-or neonatally colonized mice and PBS-inoculated controls. Splenocytes were labelled with CFSE and cultured for 4 days in the presence or absence of an *S. aureus* antigen cocktail. T cell subpopulations and proliferation were assessed by flow cytometry. Bar charts show the percentage of RORγt^+^ (Th17) cells within the total CD4^+^ T cell population (CD3^+^CD4^+^CD8^−^) or within the proliferating effector memory CD4^+^ T cell population (CFSE^low^CD3^+^CD4^+^CD8^−^CD44^+^CD62L^−^). Each dot represents pooled splenocytes from two mice (n = 12 mice/group). Data are presented as mean ± CI. Statistics: linear mixed model (including animal-ID as random factor) in two-way ANOVA design.

**S6 Fig. *S. aureus-*specific splenocytes from colonized mice secrete high amounts of IL-17A.** Splenocytes from adult-or neonatally colonized mice and PBS-inoculated controls were cultured for 4 days in the presence or absence of an *S. aureus* antigen cocktail. Cytokine concentrations were measured by bead-based multiplex assay and IFN-γ ELISA. Each dot represents pooled splenocytes from two mice (n = 12 mice/group). Data are presented as mean ± CI. Statistics: linear mixed model (including animal-ID as random factor) in two-way ANOVA design.

**S7 Fig. Early *S. aureus*-specific IFN-γ and IL-10 secretion by CLN cells following adult colonization.** Female C57BL/6NRj mice were inoculated i.n. with 1 × 10⁸ CFU *S. aureus* JSNZ or PBS. CLN cells were harvested at days 3, 6, and 10 post-colonization and cultured for 4 days in the presence or absence of an *S. aureus* antigen cocktail. Cytokine concentrations were measured by bead-based multiplex assay. Data are presented as mean ± CI (n = 8-14 mice/group). Statistics: linear mixed model (including animal-ID as random factor) in two-way ANOVA design.

**S8 Fig. Purity assessment of isolated CD4 , CD8 , and γδ T cell populations.** CLN cells from adult-or neonatally colonized mice and PBS-inoculated controls were harvested at day 28 post-colonization and pooled within each group (5 mice/group). CD4^+^, CD8^+^, and γδ T cells were isolated by magnetic bead separation. Purity was assessed by flow cytometry using Zombie NIR™ viability staining and antibodies against CD3, CD4, CD8, and TCRγδ. After doublet exclusion, live cells were gated on FSC-A/CD3, and the CD3^+^ T cell population was subsequently gated on CD4 and CD8. γδ T cells were identified within the CD4^−^CD8^−^ population by TCRγδ expression. Graphs show the percentage of each isolated population within live cells (FSC-A/CD3), within the T cell population (CD4^+^ and CD8^+^ T cells), or within the CD4^−^CD8^−^ T cell population (TCRγδ/FSC-A).

## References

1. Mulcahy ME, McLoughlin RM. Host–bacterial crosstalk determines *Staphylococcus aureus* nasal colonization. Trends in Microbiology. 2016;24: 872–886. doi:10.1016/j.tim.2016.06.012

2. Sakr A, Brégeon F, Mège J-L, Rolain J-M, Blin O. *Staphylococcus aureus* Nasal Colonization: An Update on Mechanisms, Epidemiology, Risk Factors, and Subsequent Infections. Front Microbiol. 2018;9: 2419. doi:10.3389/fmicb.2018.02419

3. Tong SYC, Davis JS, Eichenberger E, Holland TL, Fowler VG. *Staphylococcus aureus* infections: epidemiology, pathophysiology, clinical manifestations, and management. Clin Microbiol Rev. 2015;28: 603–61. doi:10.1128/CMR.00134-14

4. von Eiff C, Becker K, Machka K, Stammer H, Peters G. Nasal carriage as a source of *Staphylococcus aureus* bacteremia. Study Group. N Engl J Med. 2001;344: 11–16. doi:10.1056/NEJM200101043440102

5. Chambers HF, Fowler VG. Intertwining clonality and resistance: *Staphylococcus aureus* in the antibiotic era. J Clin Invest. 2024;134: e185824. doi:10.1172/JCI185824

6. Hajam IA, Liu GY. Linking *S. aureus* Immune Evasion Mechanisms to Staphylococcal Vaccine Failures. Antibiotics (Basel). 2024;13: 410. doi:10.3390/antibiotics13050410

7. Mrochen DM, Fernandes de Oliveira LM, Raafat D, Holtfreter S. *Staphylococcus aureus* host tropism and Its Implications for murine infection models. Int J Mol Sci. 2020;21. doi:10.3390/ijms21197061

8. Mulcahy ME, McLoughlin RM. Host-Bacterial Crosstalk Determines Staphylococcus aureus Nasal Colonization. Trends Microbiol. 2016;24: 872–886. doi:10.1016/j.tim.2016.06.012

9. Sakr A, Brégeon F, Mège J-L, Rolain J-M, Blin O. Staphylococcus aureus Nasal Colonization: An Update on Mechanisms, Epidemiology, Risk Factors, and Subsequent Infections. Front Microbiol. 2018;9. doi:10.3389/fmicb.2018.02419

10. Raineri EJM, Altulea D, van Dijl JM. Staphylococcal trafficking and infection—from ‘nose to gut’ and back. FEMS Microbiol Rev. 2022;46: fuab041. doi:10.1093/femsre/fuab041

11. Muenks CE, Hogan PG, Wang JW, Eisenstein KA, Burnham C-AD, Fritz SA. Diversity of *Staphylococcus aureus* strains colonizing various niches of the human body. J Infect. 2016;72: 698–705. doi:10.1016/j.jinf.2016.03.015

12. Lebon A, Labout JAM, Verbrugh HA, Jaddoe VWV, Hofman A, van Wamel W, et al. Dynamics and determinants of *Staphylococcus aureus* carriage in infancy: the Generation R Study. J Clin Microbiol. 2008;46: 3517–3521. doi:10.1128/JCM.00641-08

13. Peacock SJ, Justice A, Griffiths D, de Silva GDI, Kantzanou MN, Crook D, et al. Determinants of acquisition and carriage of *Staphylococcus aureus* in infancy. J Clin Microbiol. 2003;41: 5718–5725. doi:10.1128/JCM.41.12.5718-5725.2003

14. Gagnaire J, Verhoeven PO, Grattard F, Rigaill J, Lucht F, Pozzetto B, et al. Epidemiology and clinical relevance of *Staphylococcus aureus* intestinal carriage: a systematic review and meta-analysis. Expert Rev Anti Infect Ther. 2017;15: 767–785. doi:10.1080/14787210.2017.1358611

15. Matok LA, Azrad M, Leshem T, Abuzahya A, Khamaisi T, Smolkin T, et al. Mother-to-Neonate Transmission of Antibiotic-Resistant Bacteria: A Cross-Sectional Study. Microorganisms. 2021;9. doi:10.3390/microorganisms9061245

16. Scharschmidt TC. Establishing tolerance to commensal skin bacteria: timing is everything. Dermatol Clin. 2017;35: 1–9. doi:10.1016/j.det.2016.07.007

17. Olin A, Henckel E, Chen Y, Lakshmikanth T, Pou C, Mikes J, et al. Stereotypic Immune System Development in Newborn Children. Cell. 2018;174: 1277–1292. doi:10.1016/j.cell.2018.06.045

18. Piewngam P, Zheng Y, Nguyen TH, Dickey SW, Joo H-S, Villaruz AE, et al. Pathogen elimination by probiotic Bacillus via signalling interference. Nature. 2018;562: 532–537. doi:10.1038/s41586-018-0616-y

19. Esteve-Solé A, Luo Y, Vlagea A, Deyà-Martínez Á, Yagüe J, Plaza-Martín AM, et al. B Regulatory Cells: Players in Pregnancy and Early Life. Int J Mol Sci. 2018;19. doi:10.3390/ijms19072099

20. Sereme Y, Toumi E, Saifi E, Faury H, Skurnik D. Maternal immune factors involved in the prevention or facilitation of neonatal bacterial infections. Cell Immunol. 2024;395-396: 104796. doi:10.1016/j.cellimm.2023.104796

21. Archer NK, Harro JM, Shirtliff ME. Clearance of *Staphylococcus aureus* nasal carriage is T cell dependent and mediated through interleukin-17A expression and neutrophil influx. Infect Immun. 2013;81: 2070–2075. doi:10.1128/IAI.00084-13

22. Archer NK, Adappa ND, Palmer JN, Cohen NA, Harro JM, Lee SK, et al. Interleukin-17A (IL-17A) and IL-17F are critical for antimicrobial peptide production and clearance of *Staphylococcus aureus* nasal colonization. Infect Immun. 2016;84: 3575–3583. doi:10.1128/IAI.00596-16

23. Kelly AM, Leech JM, Doyle SL, McLoughlin RM. *Staphylococcus aureus*-induced immunosuppression mediated by IL-10 and IL-27 facilitates nasal colonisation. PLoS Pathog. 2022;18: e1010647. doi:10.1371/journal.ppat.1010647

24. Kolata JB, Kühbandner I, Link C, Normann N, Vu CH, Steil L, et al. The Fall of a Dogma? Unexpected High T-Cell Memory Response to *Staphylococcus aureus* in Humans. J Infect Dis. 2015;212: 830–8. doi:10.1093/infdis/jiv128

25. Zielinski CE, Mele F, Aschenbrenner D, Jarrossay D, Ronchi F, Gattorno M, et al. Pathogen-induced human TH17 cells produce IFN-γ or IL-10 and are regulated by IL-1β. Nature. 2012;484: 514–8. doi:10.1038/nature10957

26. Clegg J, Mnich ME, Carignano A, Cova G, Tavarini S, Sammicheli C, et al. *Staphylococcus aureus*-specific TIGIT+ Treg are present in the blood of healthy subjects -a hurdle for vaccination? Front Immunol. 2024;15: 1500696. doi:10.3389/fimmu.2024.1500696

27. Ferraro A, Buonocore SM, Auquier P, Nicolas I, Wallemacq H, Boutriau D, et al. Role and plasticity of Th1 and Th17 responses in immunity to *Staphylococcus aureus*. Hum Vaccin Immunother. 2019;15: 2980–2992. doi:10.1080/21645515.2019.1613126

28. Bröker B, Mrochen D, Péton V. The T cell response to *Staphylococcus aureus*. Pathogens. 2016;5: 31. doi:10.3390/pathogens5010031

29. Holtfreter S, Radcliff FJ, Grumann D, Read H, Johnson S, Monecke S, et al. Characterization of a mouse-adapted *Staphylococcus aureus* strain. Fitzgerald JR, editor. PLoS ONE. 2013;8: e71142. doi:10.1371/journal.pone.0071142

30. Mrochen DM, Grumann D, Schulz D, Gumz J, Trübe P, Pritchett-Corning K, et al. Global spread of mouse-adapted *Staphylococcus aureus* lineages CC1, CC15, and CC88 among mouse breeding facilities. Int J Med Microbiol. 2018;308: 598–606. doi:10.1016/j.ijmm.2017.11.006

31. Schulz D, Grumann D, Trübe P, Pritchett-Corning K, Johnson S, Reppschläger K, et al. Laboratory mice are frequently colonized with *Staphylococcus aureus* and mount a systemic immune response—Note of caution for In vivo infection experiments. Front Cell Infect Microbiol. 2017;7: 152. doi:10.3389/fcimb.2017.00152

32. Sun Y, Emolo C, Holtfreter S, Wiles S, Kreiswirth B, Missiakas D, et al. Staphylococcal Protein A contributes to persistent colonization of mice with *Staphylococcus aureus*. DiRita VJ, editor. J Bacteriol. 2018;200. doi:10.1128/JB.00735-17

33. Fernandes de Oliveira LM, Steindorff M, Darisipudi MN, Mrochen DM, Trübe P, Bröker BM, et al. Discovery of *Staphylococcus aureus* adhesion inhibitors by automated imaging and their characterization in a mouse model of persistent nasal colonization. Microorganisms. 2021;9. doi:10.3390/microorganisms9030631

34. Clow F, Peterken K, Pearson V, Proft T, Radcliff FJ. PilVax, a novel *Lactococcus lactis*-based mucosal vaccine platform, stimulates systemic and mucosal immune responses to *Staphylococcus aureus*. Immunol Cell Biol. 2020;98: 369–381. doi:10.1111/imcb.12325

35. Boff D, Chandrasekaran R, Putzel G, Kratofil RM, Zheng X, Castellaw A, et al. *Staphylococcus aureus* LukMF’ targets neutrophils to promote skin and soft tissue infection. Sci Adv. 2025;11: eadr5240. doi:10.1126/sciadv.adr5240

36. Langley RJ, Ting YT, Clow F, Young PG, Radcliff FJ, Choi JM, et al. Staphylococcal enterotoxin-like X (SElX) is a unique superantigen with functional features of two major families of staphylococcal virulence factors. PLoS Pathog. 2017;13: e1006549. doi:10.1371/journal.ppat.1006549

37. Xiang G, Wang Y, Ni K, Luo H, Liu Q, Song Y, et al. Nasal *Staphylococcus aureus* carriage promotes depressive behaviour in mice via sex hormone degradation. Nat Microbiol. 2025;10: 2425–2440. doi:10.1038/s41564-025-02120-6

38. Fernandes Hartzig LM, Peringathara S, Darisipudi MN, Seegert SLL, Bludau E, Weiss S, et al. Spontaneous preputial gland infection in Staphylococcus aureus-colonized male C57Bl/6 mice triggers a Th17-driven immune response. BioRxiv. 2025. 10.1101/2025.09.05.674392

39. Ge C, Monk IR, Monard SC, Bedford JG, Braverman J, Stinear TP, et al. Neutrophils play an ongoing role in preventing bacterial pneumonia by blocking the dissemination of *Staphylococcus aureus* from the upper to the lower airways. Immunol Cell Biol. 2020;98: 577–594. doi:10.1111/imcb.12343

40. Mulcahy ME, Leech JM, Renauld J-C, Mills KH, McLoughlin RM. Interleukin-22 regulates antimicrobial peptide expression and keratinocyte differentiation to control *Staphylococcus aureus* colonization of the nasal mucosa. Mucosal Immunol. 2016;9: 1429–1441. doi:10.1038/mi.2016.24

41. Cole AL, Muthukrishnan G, Chong C, Beavis A, Eade CR, Wood MP, et al. Host innate inflammatory factors and staphylococcal protein A influence the duration of human *Staphylococcus aureus* nasal carriage. Mucosal Immunol. 2016;9: 1537–48. doi:10.1038/mi.2016.2

42. Bachelerie F, Ben-Baruch A, Burkhardt AM, Combadiere C, Farber JM, Graham GJ, et al. International Union of Basic and Clinical Pharmacology. corrected. LXXXIX. Update on the extended family of chemokine receptors and introducing a new nomenclature for atypical chemokine receptors. Pharmacol Rev. 2014;66: 1–79. doi:10.1124/pr.113.007724

43. Kotpal R, S KP, Bhalla P, Dewan R, Kaur R. Incidence and Risk Factors of Nasal Carriage of *Staphylococcus aureus* in HIV-Infected Individuals in Comparison to HIV-Uninfected Individuals: A Case-Control Study. J Int Assoc Provid AIDS Care. 2016;15: 141–7. doi:10.1177/2325957414554005

44. Nurjadi D, Kain M, Marcinek P, Gaile M, Heeg K, Zanger P. Ratio of T-Helper Type 1 (Th1) to Th17 Cytokines in Whole Blood Is Associated With Human β-Defensin 3 Expression in Skin and Persistent *Staphylococcus aureus* Nasal Carriage. J Infect Dis. 2016;214: 1744–1751. doi:10.1093/infdis/jiw440

45. Reiss-Mandel A, Rubin C, Zayoud M, Rahav G, Regev-Yochay G. *Staphylococcus aureus* Colonization Induces Strain-Specific Suppression of Interleukin-17. Torres VJ, editor. Infect Immun. 2018;86: e00834–17. doi:10.1128/IAI.00834-17

46. Zhang F, Ledue O, Jun M, Goulart C, Malley R, Lu Y-J. Protection against *Staphylococcus aureus* Colonization and Infection by B-and T-Cell-Mediated Mechanisms. mBio. 2018;9. doi:10.1128/mBio.01949-18

47. Wertheim HFL, Vos MC, Ott A, van Belkum A, Voss A, Kluytmans JAJW, et al. Risk and outcome of nosocomial *Staphylococcus aureus* bacteraemia in nasal carriers versus non-carriers. Lancet. 2004;364: 703–5. doi:10.1016/S0140-6736(04)16897-9

48. Wertheim HFL, Melles DC, Vos MC, van Leeuwen W, van Belkum A, Verbrugh HA, et al. The role of nasal carriage in *Staphylococcus aureus* infections. The Lancet Infectious Diseases. 2005;5: 751–62. doi:10.1016/S1473-3099(05)70295-4

49. Wymore Brand M, Proctor AL, Hostetter JM, Zhou N, Friedberg I, Jergens AE, et al. Vertical transmission of attaching and invasive *E. coli* from the dam to neonatal mice predisposes to more severe colitis following exposure to a colitic insult later in life. Nakano H, editor. PLoS ONE. 2022;17: e0266005. doi:10.1371/journal.pone.0266005

50. Andrade EB, Magalhães A, Puga A, Costa M, Bravo J, Portugal CC, et al. A mouse model reproducing the pathophysiology of neonatal group B streptococcal infection. Nat Commun. 2018;9: 3138. doi:10.1038/s41467-018-05492-y

51. Flaxman A, Yamaguchi Y, van Diemen PM, Rollier C, Allen E, Elshina E, et al. Heterogeneous early immune responses to the *S. aureus* EapH2 antigen induced by gastrointestinal tract colonisation impact the response to subsequent vaccination. Vaccine. 2019;37: 494–501. doi:10.1016/j.vaccine.2018.11.063

52. Flaxman A, van Diemen PM, Yamaguchi Y, Allen E, Lindemann C, Rollier CS, et al. Development of persistent gastrointestinal *S. aureus* carriage in mice. Sci Rep. 2017;7: 12415. doi:10.1038/s41598-017-12576-0

53. Iwase T, Uehara Y, Shinji H, Tajima A, Seo H, Takada K, et al. *Staphylococcus epidermidis* Esp inhibits *Staphylococcus aureus* biofilm formation and nasal colonization. Nature. 2010;465: 346–9. doi:10.1038/nature09074

54. Zipperer A, Konnerth MC, Laux C, Berscheid A, Janek D, Weidenmaier C, et al. Human commensals producing a novel antibiotic impair pathogen colonization. Nature. 2016;535: 511–516. doi:10.1038/nature18634

55. Laux C, Peschel A, Krismer B. *Staphylococcus aureus* Colonization of the Human Nose and Interaction with Other Microbiome Members. Microbiol Spectr. 2019;7. doi:10.1128/microbiolspec.GPP3-0029-2018

56. Scharschmidt TC, Vasquez KS, Truong H-A, Gearty SV, Pauli ML, Nosbaum A, et al. A wave of regulatory T cells into neonatal skin mediates tolerance to commensal microbes. Immunity. 2015;43: 1011–21. doi:10.1016/j.immuni.2015.10.016

57. Mork RL, Hogan PG, Muenks CE, Boyle MG, Thompson RM, Sullivan ML, et al. Longitudinal, strain-specific *Staphylococcus aureus* introduction and transmission events in households of children with community-associated meticillin-resistant *S. aureus* skin and soft tissue infection: a prospective cohort study. Lancet Infect Dis. 2020;20: 188–198. doi:10.1016/S1473-3099(19)30570-5

58. Satorres SE, Alcaráz LE, Cargnelutti E, Di Genaro MS. IFN-γ plays a detrimental role in murine defense against nasal colonization of *Staphylococcus aureus*. Immunol Lett. 2009;123: 185–188. doi:10.1016/j.imlet.2009.03.003

59. Montgomery CP, Daniels M, Zhao F, Alegre M-L, Chong AS, Daum RS. Protective Immunity against Recurrent *Staphylococcus aureus* Skin Infection Requires Antibody and Interleukin-17A. Camilli A, editor. Infect Immun. 2014;82: 2125–2134. doi:10.1128/IAI.01491-14

60. Köckritz-Blickwede M, Rohde M, Oehmcke S, Miller LS, Cheung AL, Herwald H, et al. Immunological mechanisms underlying the genetic predisposition to severe *Staphylococcus aureus* infection in the mouse model. Am J Pathol. 2008;173: 1657–68. doi:10.2353/ajpath.2008.080337

61. Lejeune A, Zhou C, Ercelen D, Putzel G, Yao X, Guy AR, et al. Sex-dependent gastrointestinal colonization resistance to MRSA is microbiota and Th17 dependent. eLife. 2025;13: RP101606. doi:10.7554/eLife.101606

62. Bogaert D, Weinberger D, Thompson C, Lipsitch M, Malley R. Impaired innate and adaptive immunity to *Streptococcus pneumoniae* and its effect on colonization in an infant mouse model. Infect Immun. 2009;77: 1613–22. doi:10.1128/IAI.00871-08

63. Zhang Q, Coveney AP, Yu S, Liu JH, Li Y, Blankson S, et al. Inefficient antimicrobial functions of innate phagocytes render infant mice more susceptible to bacterial infection. Eur J Immunol. 2013;43: 1322–32. doi:10.1002/eji.201243077

64. Wolfgramm H, Busch LM, Tebben J, Mehlan H, Hagenau L, Sura T, et al. Integrated genomic and proteomic analysis of the mouse-adapted *Staphylococcus aureus* strain JSNZ. Curr Res Microb Sci. 2025;9: 100489. doi:10.1016/j.crmicr.2025.100489

65. Braun C, Badiou C, Guironnet-Paquet A, Iwata M, Lenief V, Mosnier A, et al. *Staphylococcus aureus*-specific skin resident memory T cells protect against bacteria colonization but exacerbate atopic dermatitis-like flares in mice. J Allergy Clin Immunol. 2024;154: 355–374. doi:10.1016/j.jaci.2024.03.032

66. Xu R, Shears RK, Sharma R, Krishna M, Webb C, Ali R, et al. IL-35 is critical in suppressing superantigenic *Staphylococcus aureus*-driven inflammatory Th17 responses in human nasopharynx-associated lymphoid tissue. Mucosal Immunol. 2020;13: 460–470. doi:10.1038/s41385-019-0246-1

67. Murphy AG, O’Keeffe KM, Lalor SJ, Maher BM, Mills KHG, McLoughlin RM. *Staphylococcus aureus* infection of mice expands a population of memory γδ T cells that are protective against subsequent infection. J Immunol. 2014;192: 3697–708. doi:10.4049/jimmunol.1303420

68. Cho JS, Pietras EM, Garcia NC, Ramos RI, Farzam DM, Monroe HR, et al. IL-17 is essential for host defense against cutaneous *Staphylococcus aureus* infection in mice. J Clin Invest. 2010;120: 1762–73. doi:10.1172/JCI40891

69. Marchitto MC, Dillen CA, Liu H, Miller RJ, Archer NK, Ortines RV, et al. Clonal Vγ6^+^ Vδ4^+^ T cells promote IL-17–mediated immunity against *Staphylococcus aureus* skin infection. Proc Natl Acad Sci USA. 2019;116: 10917–10926. doi:10.1073/pnas.1818256116

70. Kelly AM, McCarthy KN, Claxton TJ, Carlile SR, O’Brien EC, Vozza EG, et al. IL-10 inhibition during immunization improves vaccine-induced protection against *Staphylococcus aureus* infection. JCI Insight. 2024;9: e178216. doi:10.1172/jci.insight.178216

71. Bonifacius A, Goldmann O, Floess S, Holtfreter S, Robert PA, Nordengrün M, et al. *Staphylococcus aureus* alpha-toxin limits type 1 while fostering type 3 immune responses. Front Immunol. 2020;11: 1579. doi:10.3389/fimmu.2020.01579

72. López-Yglesias AH, Burger E, Araujo A, Martin AT, Yarovinsky F. T-bet-independent Th1 response induces intestinal immunopathology during *Toxoplasma gondii* infection. Mucosal Immunol. 2018;11: 921–931. doi:10.1038/mi.2017.102

73. Uebele J, Stein C, Nguyen M-T, Schneider A, Kleinert F, Tichá O, et al. Antigen delivery to dendritic cells shapes human CD4+ and CD8+ T cell memory responses to *Staphylococcus aureus*. Miller LS, editor. PLoS Pathog. 2017;13: e1006387. doi:10.1371/journal.ppat.1006387

74. Stentzel S, Teufelberger A, Nordengrün M, Kolata J, Schmidt F, van Crombruggen K, et al. Staphylococcal serine protease-like proteins are pacemakers of allergic airway reactions to *Staphylococcus aureus*. J Allergy Clin Immunol. 2017;139: 492–500. doi:10.1016/j.jaci.2016.03.045

75. Nordengrün M, Abdurrahman G, Treffon J, Wächter H, Kahl BC, Bröker BM. Allergic Reactions to Serine Protease-Like Proteins of *Staphylococcus aureus*. Front Immunol. 2021;12: 651060. doi:10.3389/fimmu.2021.651060

76. Teufelberger AR, Bröker BM, Krysko DV, Bachert C, Krysko O. *Staphylococcus aureus* orchestrates type 2 airway diseases. Trends Mol Med. 2019;25: 696–707. doi:10.1016/j.molmed.2019.05.003

77. Leech JM, Lacey KA, Mulcahy ME, Medina E, McLoughlin RM. IL-10 plays opposing roles during *Staphylococcus aureus* systemic and localized infections. J Immunol. 2017;198: 2352–2365. doi:10.4049/jimmunol.1601018

78. Heim CE, Vidlak D, Kielian T. Interleukin-10 production by myeloid-derived suppressor cells contributes to bacterial persistence during *Staphylococcus aureus* orthopedic biofilm infection. J Leukoc Biol. 2015;98: 1003–13. doi:10.1189/jlb.4VMA0315-125RR

79. Sanchez M, Kolar SL, Müller S, Reyes CN, Wolf AJ, Ogawa C, et al. O-Acetylation of peptidoglycan limits Helper T Cell priming and permits *Staphylococcus aureus* reinfection. Cell Host Microbe. 2017;22: 543–551. doi:10.1016/j.chom.2017.08.008

80. Rasquinha MT, Sur M, Lasrado N, Reddy J. IL-10 as a Th2 cytokine: Differences between mice and humans. J Immunol. 2021;207: 2205–2215. doi:10.4049/jimmunol.2100565

81. Frodermann V, Chau TA, Sayedyahossein S, Toth JM, Heinrichs DE, Madrenas J. A modulatory interleukin-10 response to staphylococcal peptidoglycan prevents Th1/Th17 adaptive immunity to *Staphylococcus aureus*. J Infect Dis. 2011;204: 253–62. doi:10.1093/infdis/jir276

82. Enroth TJ, Severn MM, Costa FG, Bovee AR, Wilkening RV, Nguyen DT, et al. Global changes in *Staphylococcus aureus* virulence and metabolism during colonization of healthy skin. Torres VJ, editor. Infect Immun. 2025;93: e00028–25. doi:10.1128/iai.00028-25

83. Brown AF, Leech JM, Rogers TR, McLoughlin RM. *Staphylococcus aureus* colonization: Modulation of host immune response and impact on human vaccine design. Front Immunol. 2014;4: 20. doi:10.3389/fimmu.2013.00507

84. Mazmanian SK, Liu CH, Tzianabos AO, Kasper DL. An immunomodulatory molecule of symbiotic bacteria directs maturation of the host immune system. Cell. 2005;122: 107–118. doi:10.1016/j.cell.2005.05.007

85. Gensollen T, Iyer SS, Kasper DL, Blumberg RS. How colonization by microbiota in early life shapes the immune system. Science. 2016;352: 539–544. doi:10.1126/science.aad9378

86. Atarashi K, Tanoue T, Shima T, Imaoka A, Kuwahara T, Momose Y, et al. Induction of colonic regulatory T cells by indigenous Clostridium species. Science. 2011;331: 337–341. doi:10.1126/science.1198469

87. Leech JM, Dhariwala MO, Lowe MM, Chu K, Merana GR, Cornuot C, et al. Toxin-triggered Interleukin-1 receptor signaling enables early-life discrimination of pathogenic versus commensal skin bacteria. Cell Host Microbe. 2019;26: 795–809. doi:10.1016/j.chom.2019.10.007

88. Fitzpatrick EA, You D, Shrestha B, Siefker D, Patel VS, Yadav N, et al. A neonatal murine model of MRSA pneumonia. PLoS ONE. 2017;12: 0169273. doi:10.1371/journal.pone.0169273

89. Tebartz C, Horst SA, Sparwasser T, Huehn J, Beineke A, Peters G, et al. A major role for myeloid-derived suppressor cells and a minor role for regulatory T cells in immunosuppression during *Staphylococcus aureus* infection. J Immunol. 2015;194: 1100–11. doi:10.4049/jimmunol.1400196

90. Holtfreter S, Kolata J, Stentzel S, Bauerfeind S, Schmidt F, Sundaramoorthy N, et al. Omics Approaches for the Study of Adaptive Immunity to *Staphylococcus aureus* and the Selection of Vaccine Candidates. Proteomes. 2016;4. doi:10.3390/proteomes4010011

91. R Core Team. R: A Language and Environment for Statistical Computing. Vienna, Austria: R Foundation for Statistical Computing; 2025. Available: https://www.R-project.org/

92. Wickham H, Averick M, Bryan J, Chang W, McGowan L, François R, et al. Welcome to the Tidyverse. JOSS. 2019;4: 1686. doi:10.21105/joss.01686

93. Kassambara A. rstatix: Pipe-Friendly Framework for Basic Statistical Tests. R package version 0.7.2. 1 Jan 2023.

94. Wickham, Hadley. ggplot2: Elegant Graphics for Data Analysis. New York: Springer-Verlag; 2016. Available: https://ggplot2-book.org/

