## Supplementary figures and images for "A mouse-adapted *Staphylococcus aureus* strain enables lifelong neonatal colonization and elicits a Th17-dominated immune response"

### Supplemental Figures 1-8

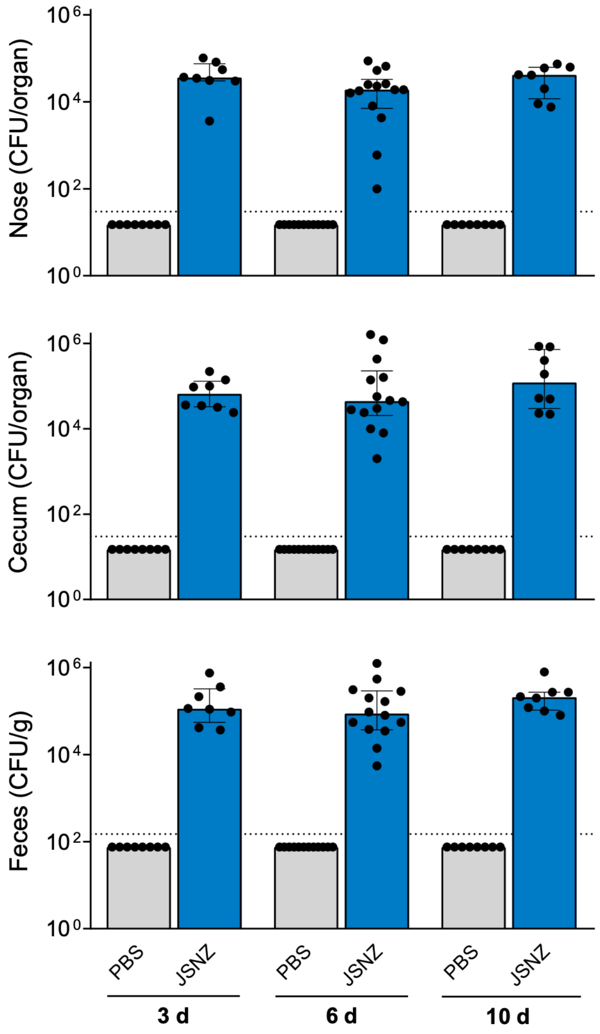

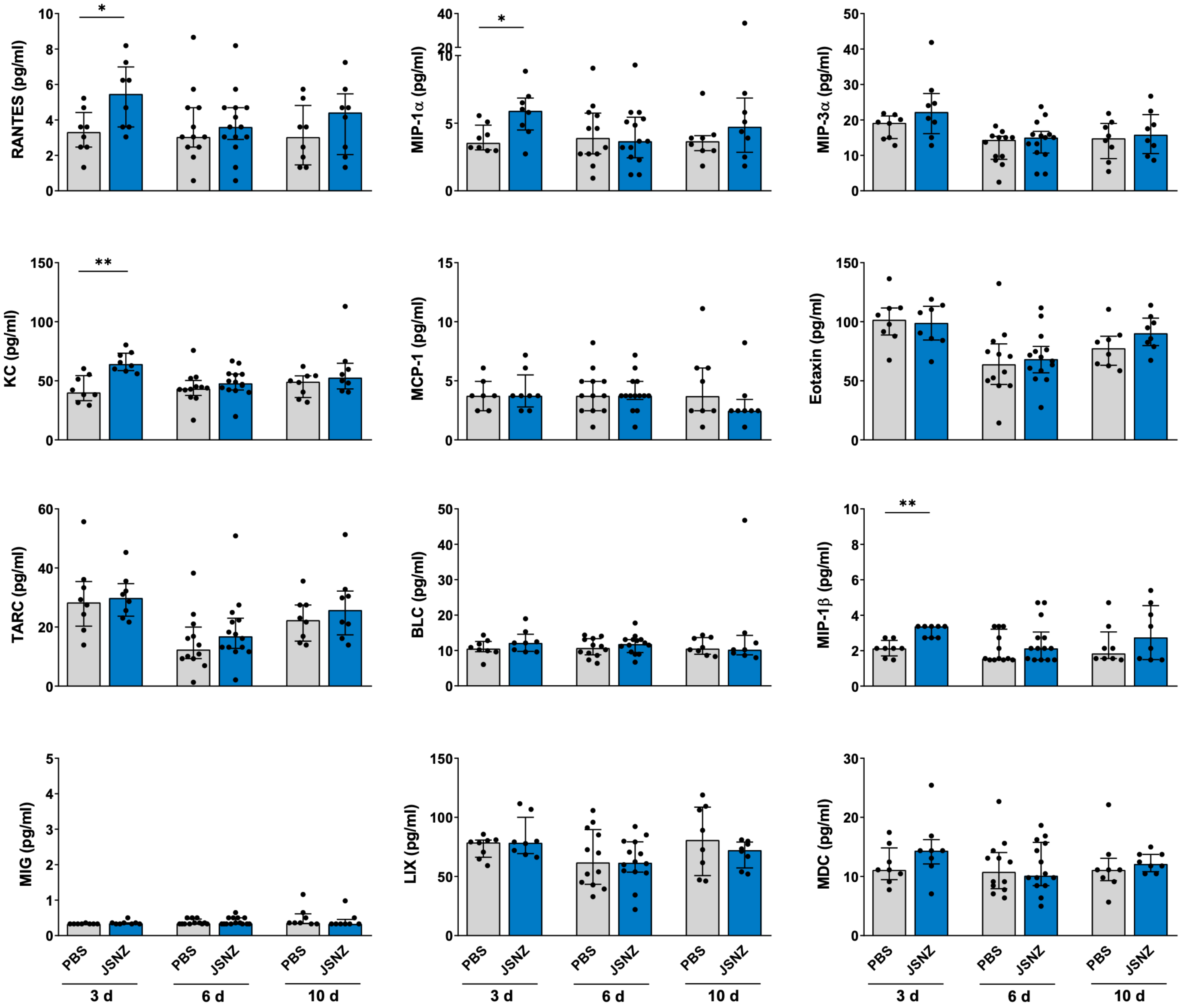

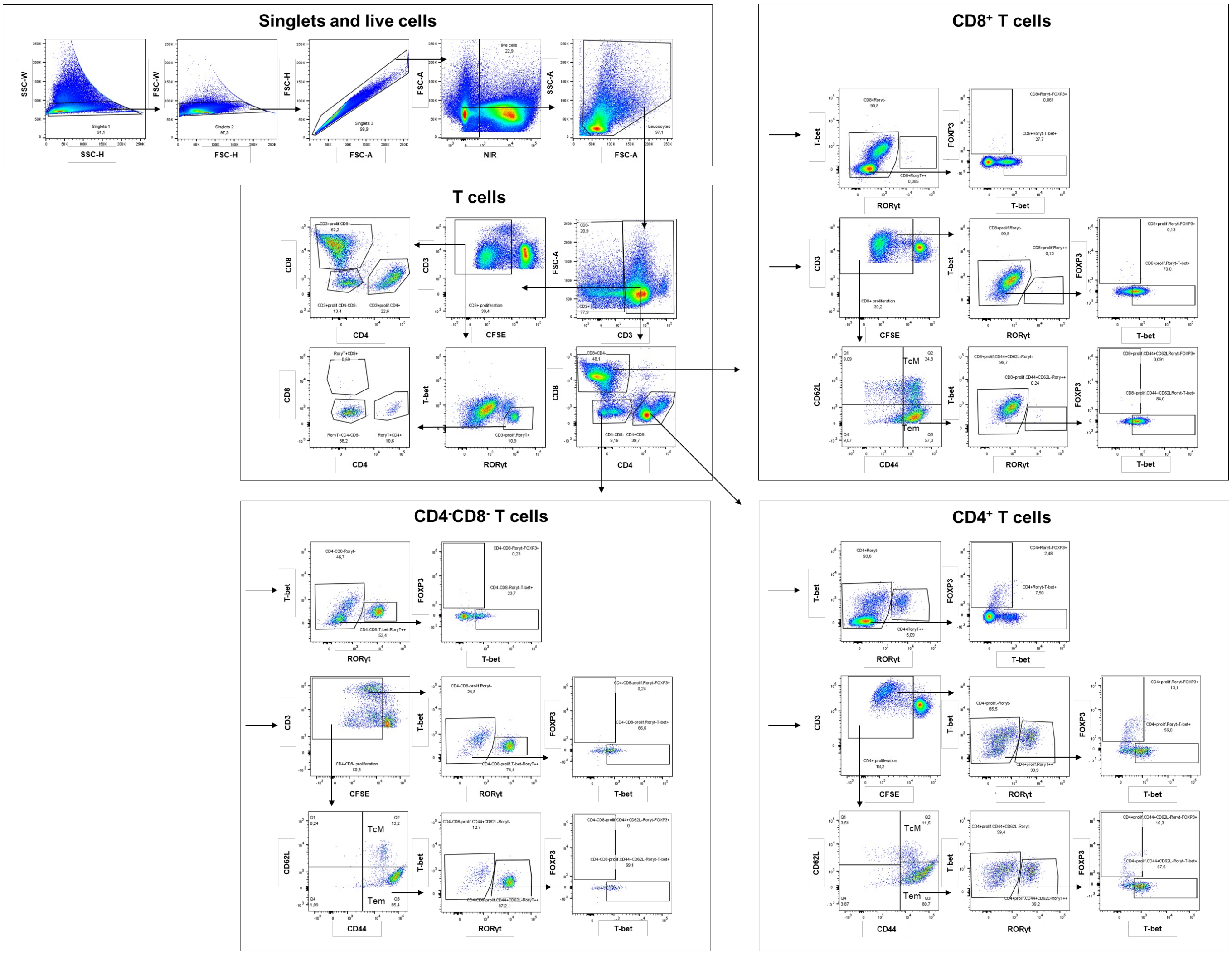

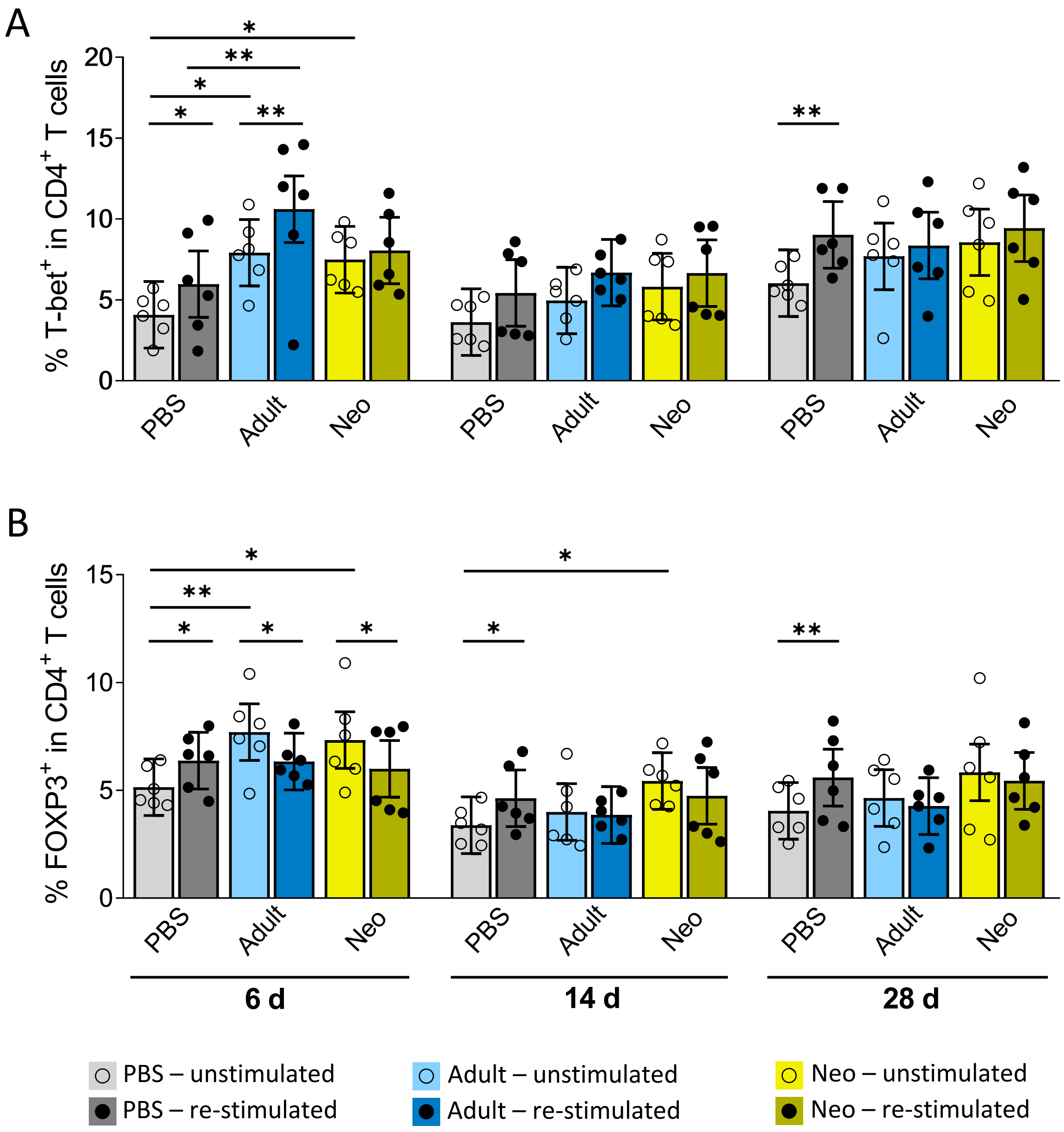

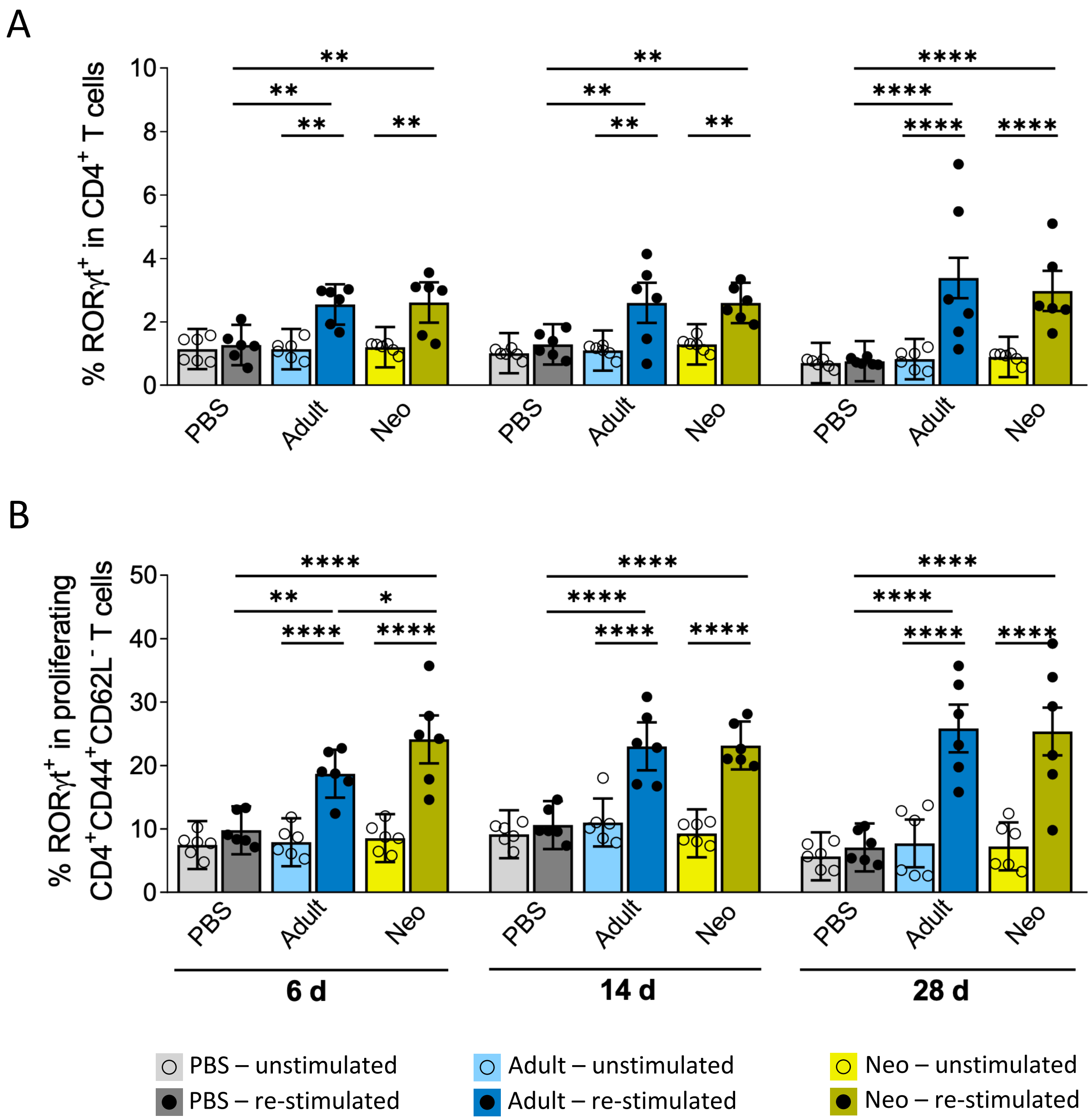

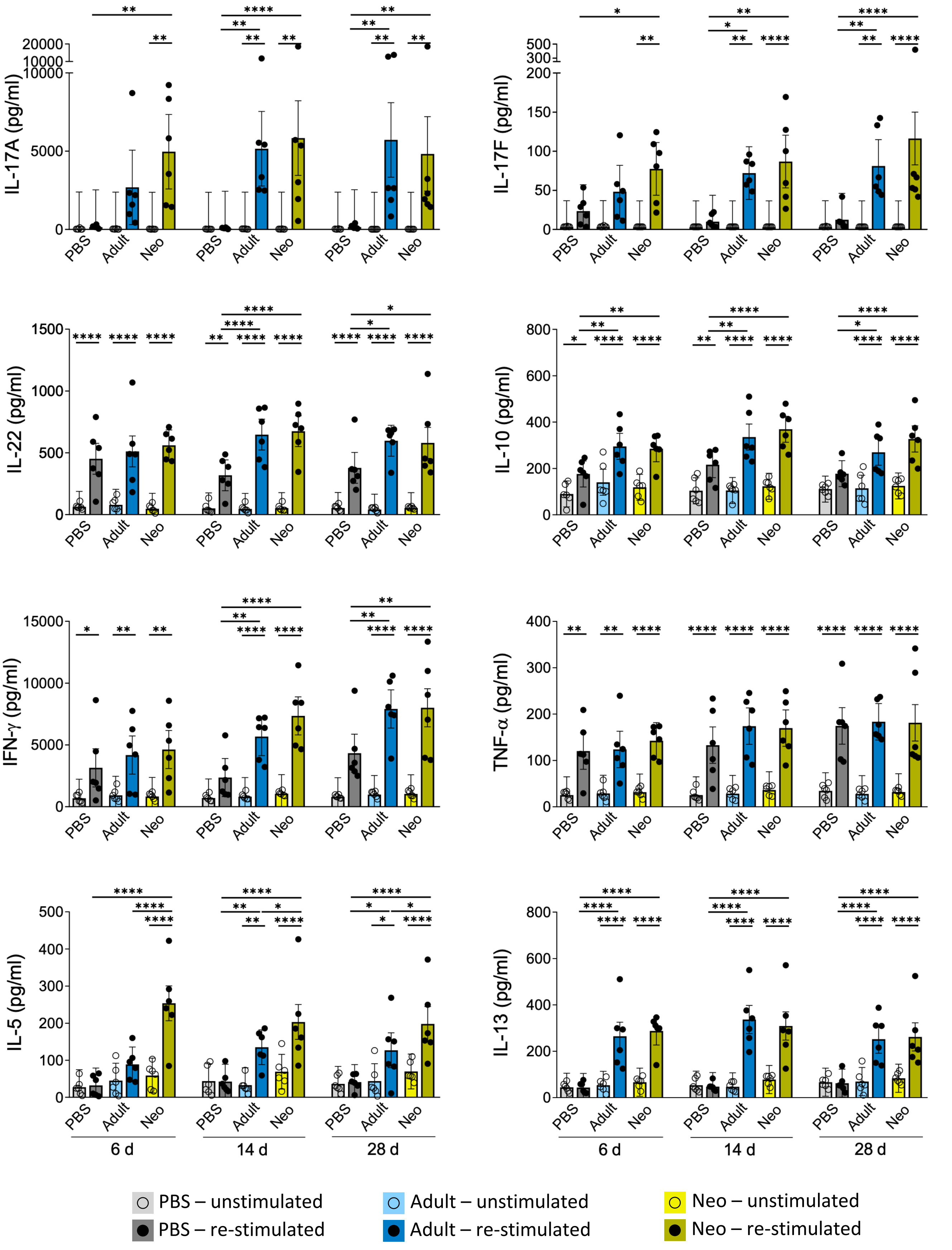

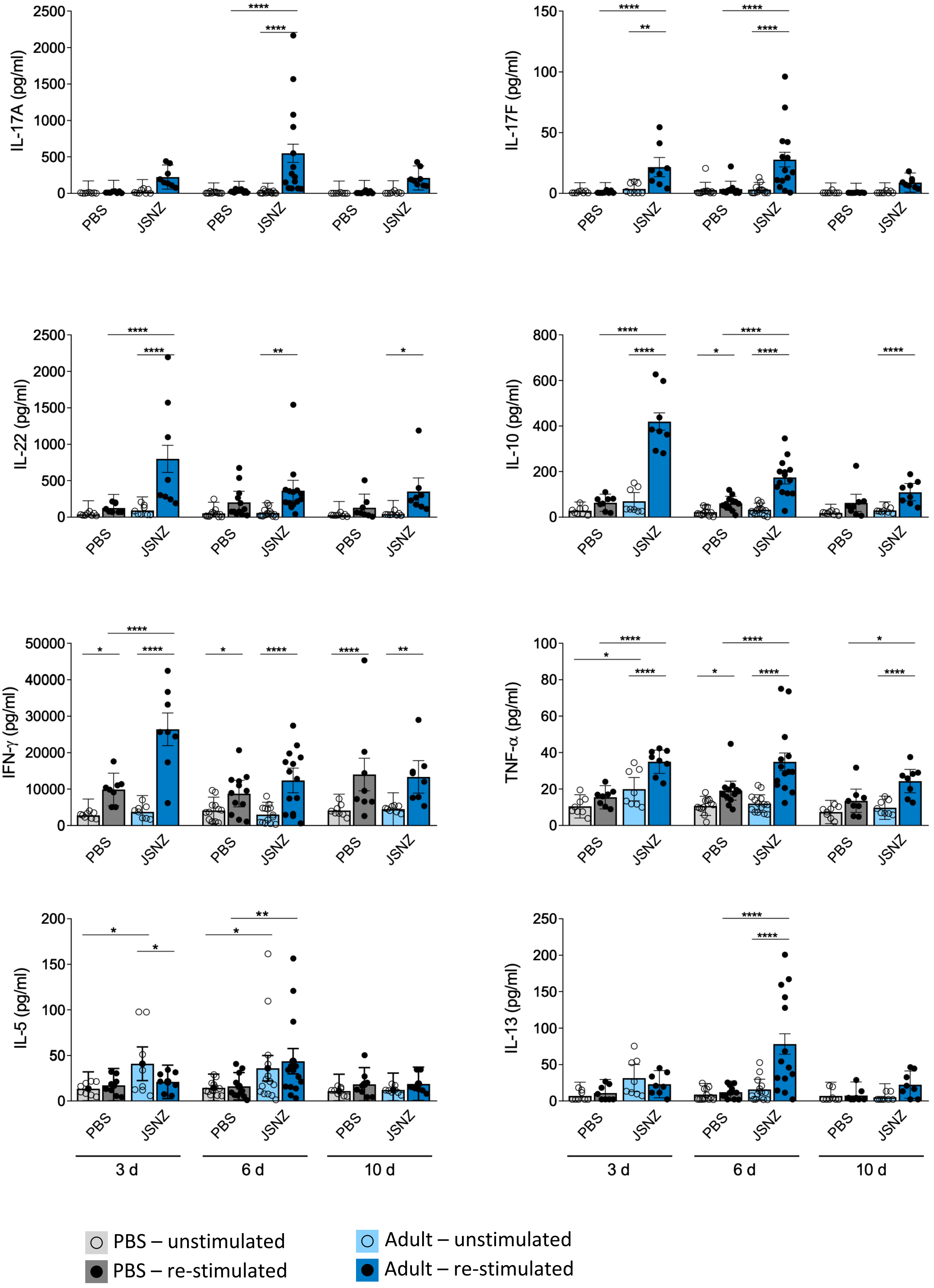

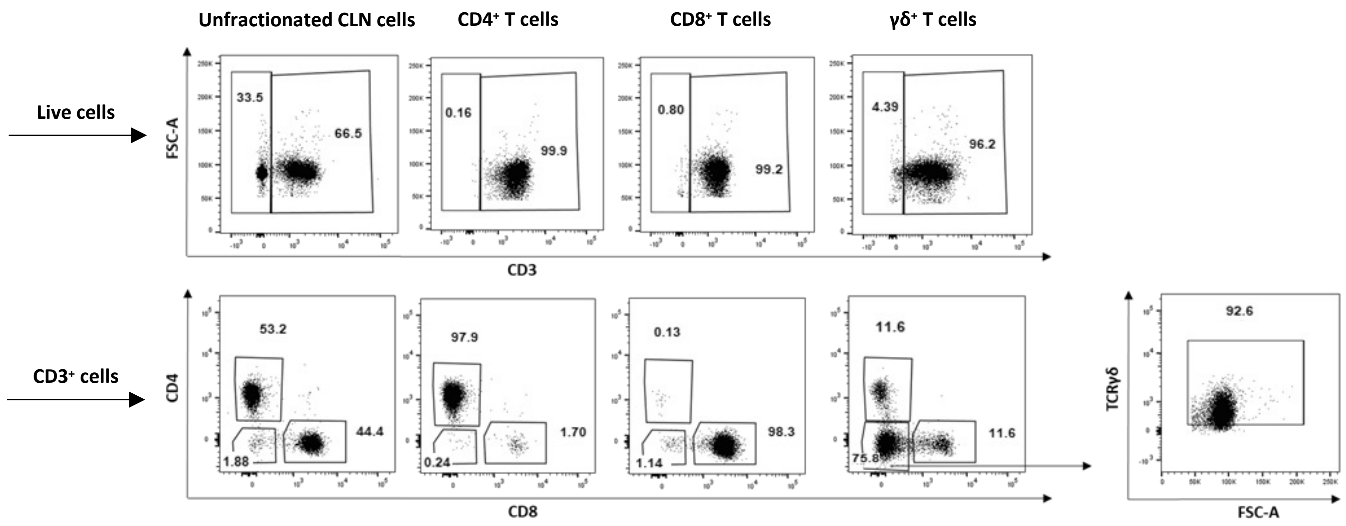
