## Supplemental Tables 1-2 for "A mouse-adapted *Staphylococcus aureus* strain enables lifelong neonatal colonization and elicits a Th17-dominated immune response"

S1 Table. Antibodies used for flow cytometry of T cell proliferation

| **Marker**  **(Anti-mouse antibodies)** | **Conjugate** | **Clone** | **Dilution** | **Provider** | **Catalog** |
| --- | --- | --- | --- | --- | --- |
| CD3 | BV650 | 17A2 | 1:10 | Biolegend | 100229 |
| CD4 | BV605 | RM4-5 | 1:100 | Biolegend | 100548 |
| CD8 | BV510 | 53-6.7 | 1:20 | Biolegend | 100752 |
| CD62L | PerCPCy5.5 | MEL-14 | 1:40 | Biolegend | 104432 |
| CD44 | Pe-Cy7 | IM7 | 1:40 | Biolegend | 103030 |
| RORγt | PE | Q31-378 | 1:50 | BD Biosciences | 562607 |
| FoxP3 | VioR667 | REA788 | 1:50 | Miltenyi Biotec | 130111604 |
| T-bet | BV421 | 4B10 | 1:25 | Biolegend | 644816 |

S2 Table. Antibodies used for flow cytometry of isolated T cells

| **Marker**  **(Anti-mouse antibodies)** | **Conjugate** | **Clone** | **Dilution** | **Provider** | **Catalog** |
| --- | --- | --- | --- | --- | --- |
| CD3 | PerCP-Vio700 | Rea641 | 1:50 | Miltenyi Biotec | 130120826 |
| CD4 | Vio R720 | Rea604 | 1:50 | Miltenyi Biotec | 130127473 |
| CD8 | VioGreen | Rea601 | 1:50 | Miltenyi Biotec | 130122017 |
| TCRγ/δ | PE-Vio770 | Rea633 | 1:50 | Miltenyi Biotec | 130123290 |
